# Subtype-specific and co-occurring genetic alterations in B-cell non-Hodgkin lymphoma

**DOI:** 10.1101/674259

**Authors:** Man Chun John Ma, Saber Tadros, Alyssa Bouska, Tayla Heavican, Haopeng Yang, Qing Deng, Dalia Moore, Ariz Akhter, Keenan Hartert, Neeraj Jain, Jordan Showell, Sreejoyee Ghosh, Lesley Street, Marta Davidson, Christopher Carey, Joshua Tobin, Deepak Perumal, Julie M. Vose, Matthew A. Lunning, Aliyah R. Sohani, Benjamin J. Chen, Shannon Buckley, Loretta J. Nastoupil, R. Eric Davis, Jason R. Westin, Nathan H. Fowler, Samir Parekh, Maher Gandhi, Sattva Neelapu, Douglas Stewart, Kapil Bhalla, Javeed Iqbal, Timothy Greiner, Scott J. Rodig, Adnan Mansoor, Michael R. Green

## Abstract

B-cell non-Hodgkin’s lymphoma (B-NHL) encompasses multiple clinically and phenotypically distinct subtypes of malignancy with unique molecular etiologies. Common subtypes of B-NHL such as diffuse large B-cell lymphoma (DLBCL) have been comprehensively interrogated at the genomic level. But rarer subtypes such as mantle cell lymphoma (MCL) remain sparsely characterized. Furthermore, multiple B-NHL subtypes have thus far not been comprehensively compared using the same methodology to identify conserved or subtype-specific patterns of genomic alterations. Here, we employed a large targeted hybrid-capture sequencing approach encompassing 380 genes to interrogate the genomic landscapes of 685 B-NHL tumors at high depth; including DLBCL, MCL, follicular lymphoma (FL), and Burkitt lymphoma (BL). We identified conserved hallmarks of B-NHL that were deregulated in the majority of tumor from each subtype, including the frequent genetic deregulation of the ubiquitin proteasome system (UPS). In addition, we identified subtype-specific patterns of genetic alterations, including clusters of co-occurring mutations and DNA copy number alterations. The cumulative burden of mutations within a single cluster were more discriminatory of B-NHL subtypes than individual mutations, implicating likely patterns of genetic cooperation that contribute to disease etiology. We therefore provide the first cross-sectional analysis of mutations and DNA copy number alterations across major B-NHL subtypes and a framework of co-occurring genetic alterations that deregulate genetic hallmarks and likely cooperate in lymphomagenesis.

## INTRODUCTION

Non-Hodgkin’s lymphomas (NHL) are a heterogeneous group of lymphoid malignancies that predominantly arise from mature B-cells (B-NHL). Although mature B-cell neoplasms encompass 38 unique diagnostic subtypes, over 85% of cases fall within only 7 histologies(1, 2). Recent next generation sequencing (NGS) studies have shed light onto the key driver mutations in many of these B-NHL subtypes; for example, large studies of diffuse large B-cell lymphoma (DLBCL) have led to proposed genomic subtypes that have unique etiologies(3-5). However, many less common NHL subtypes such as mantle cell lymphoma (MCL) have not been as extensively characterized(6, 7). Furthermore, until recently(3, 4) genetic alterations have been considered in a binary fashion as either driver events, which directly promote disease genesis or progression, or passenger events, which have little/no impact on disease biology. In contrast to this principal, most B-NHLs do not result from a single dominant driver but instead result from the serial acquisition of genetic alterations that cooperate in lymphomagenesis(8). The genetic context of each mutation likely determines its oncogenic potential, and groups of mutations should therefore be considered collectively rather than on as singular events. For example, the ‘C5’ and ‘MCD’ clusters identified in DLBCL by Chapuy *et al*. and Schmitz *et al*., respectively, are characterized by the co-occurrence of *CD79B* and *MYD88* mutations(3, 4). In animal models, the *Myd88* L252P mutation (equivalent to human L265P) was found to promote down-regulation of surface IgM and a phenotype resembling B-cell anergy(9). However, this effect could be rescued by *Cd79b* mutation, showing that these co-occurring mutations cooperate(9). The characterization of other significantly co-occurring genetic alterations are therefore likely to reveal additional important cooperative relationships. We approached this challenge by performing genomic profiling of 685 B-NHLs across different subtypes. Through this cross-sectional analysis we characterized genomic hallmarks of B-NHL and sets of significantly co-associated events that likely represent subtype-specific cooperating genetic alterations. This study therefore provides new insight into how co-occurring clusters of genetic alterations may contribute to molecularly and phenotypically distinct subtypes of B-NHL.

## METHODS

An overview of our approach is shown in Figure S1. For detailed methods, please refer to supplementary information.

### Tumor DNA samples

We collected DNA from 685 B-NHL tumors, including 199 follicular lymphoma (FL), 196 mantle cell lymphoma (MCL), 148 diffuse large B-cell lymphoma (DLBCL), 107 Burkitt’s lymphoma (BL), 21 high-grade B-cell lymphoma not otherwise specified (HGBL-NOS), and 14 high-grade B-cell lymphoma with *MYC, BCL2* and/or *BCL6* rearrangement (DHL) (Table S1). A total of 462 samples were obtained from the University of Nebraska Medical Center, and were prioritized for inclusion in this study if they had been previously undergone pathology review and been interrogated by Affymetrix U133 Plus 2.0 gene expression microarrays(10-12) (n=284). An additional series of 223 FFPE tumors were collected from other centers. Samples were de-identified and accompanied by the diagnosis from the medical records, plus overall survival time and status when available. Medical record diagnosis was used in all cases except for those with fluorescence *in situ* hybridization showing translocations in *MYC* and *BCL2* and/or *BCL6*, which were amended to DHL. Sequencing results for a subset of 52 BL tumors were described previously(13). All MCL samples were either positive for *CCND1* translocation by FISH or positive for CCND1 protein expression by immunohistochemistry, depending on the diagnostic practices of the contributing institution.

### Next generation sequencing

A total of 500-1000ng of gDNA was sonicated using a Covaris S2 Ultrasonicator, and libraries prepared using KAPA Hyper Prep Kits (Roche) with TruSeq Adapters (Bioo Scientific) and a maximum of 8 cycles of PCR (average of 4 cycles). Libraries were qualified by TapeStation 4200, quantified by Qubit and 10-to 12-plexed for hybrid capture. Each multiplexed library was enriched using our custom LymphoSeq panel encompassing the full coding sequences of 380 genes that were determined to be somatically mutated in B-cell lymphoma (Table S2, Supplementary Methods), as well as tiling recurrent translocation breakpoints. Enrichments were amplified with 4-8 cycles of PCR and sequenced on a single lane of an Illumina HiSeq 4000 with 100PE reads in high-output mode at the Hudson Alpha Institute for Biotechnology or the MD Anderson Sequencing and Microarray Facility. Variants were called using our previously validated ensemble approach(13, 14), germline polymorphisms were filtered using dbSNP annotation and the EXAC dataset containing 60,706 healthy individuals(15), and significantly mutated genes were defined by MutSig2CV(16). Copy number alterations identified by CopyWriteR(17), which was validated using 3 FL tumors with matched Affymetrix 250K SNP array (Figure S2), and significant DNA copy number alterations were determined by GISTIC2(18). Translocations were called using FACTERA(19), which we previously validated against MYC translocation status determined by FISH(20). Mutation and CNA data are publicly viewable through cBioPortal: https://www.cbioportal.org/study/summary?id=mbn_mdacc_2013. Matched gene expression microarray data are available through the Gene Expression Omnibus, accession GSE132929. For further details, refer to supplementary methods.

## RESULTS

### Identification of significantly mutated genes and structural alterations

We used a 380 gene custom targeted sequencing approach, LymphoSeq, to interrogate the genomes of 685 mature B-NHLs, sequencing to an average depth of 578X (Min, 101X; Max, 1785X; Table S1) for a total yield of 1.81 Tbp. Somatic nucleotide variants (SNVs) and small insertions/deletions (InDels) were identified using an ensemble approach that we have previously validated(14) (Table S3) and significantly mutated genes were identified using MutSig2CV (Table S4). Matched germline DNA was available from purified T-cells of 20 tumors (11 FL and 9 MCL) and sequenced to validate the filtering of germline variants; 0/632 variants called within these tumors were identified in the matched germline samples, which indicates that the filtering of germline variants was effective. Genes that were significantly mutated in the full cohort or in any one of the 4 subtypes with >100 tumors (BL, DLBCL, FL, MCL) were included, as well as frequently mutated genes that are targets of AID (Table S5, **Figure 1**). Predictably, AID-associated mutations were higher in frequency among GC-derived lymphomas (BL, DLBCL, FL), but also accounted for 7.6% of all coding and non-coding mutations in MCL (Table S6). The mutational burden calculated from our targeted region significantly correlated with that from the whole exome (Figure S3A) and was significantly higher in DLBCL and other high-grade tumors compared to FL and MCL (**Figure 1**; Figure S3B).

**Figure 1:**
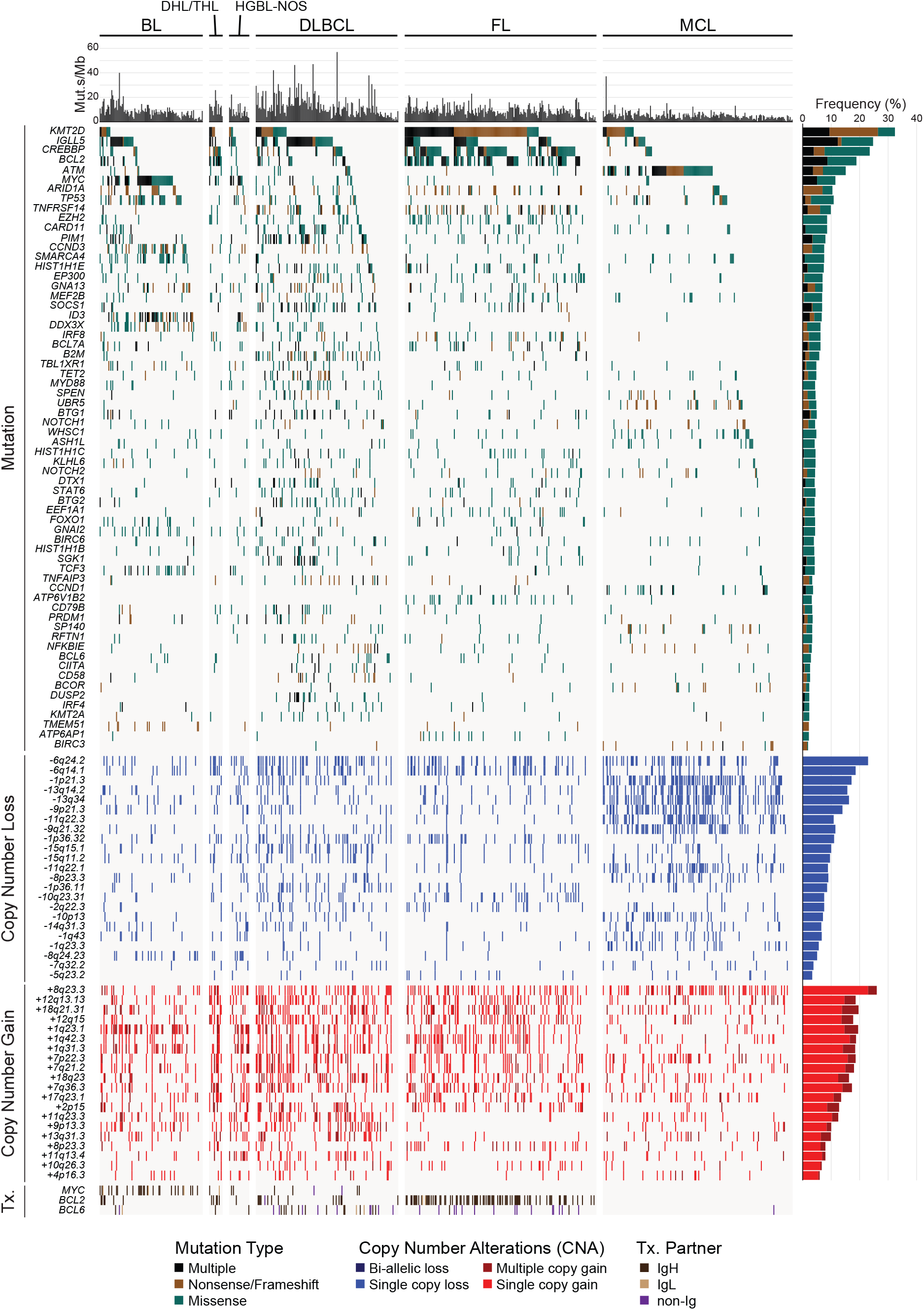
Recurrently mutated genes in B-NHL subtypes. An oncoplot shows significantly mutated genes, DNA copy number alterations and translocations (Tx.) across our cohort of 685 B-NHL tumors. Mutation types and frequencies are summarized for each gene/CNA on the right, and the mutational burden for each case are shown at the top.

The hybrid capture probes utilized in our design also targeted recurrent breakpoint regions in the immunoglobulin heavy- and light-chain loci, and recurrent breakpoints in or near the *BCL2, MYC* and *BCL6* genes, and translocations were called using a method that detects discordantly mapped reads(19) (**Figures 1 and 2A**). Our prior validation of this approach in cases with matched fluorescence in situ hybridization (FISH) data for *MYC* showed that it is 100% specific, but only ∼40% sensitive for translocation detection(13). This limit of sensitivity likely varies for different genes depending on how well the breakpoints are clustered into hotspots that are targeted by our capture probes. Nonetheless, we observed a significantly higher fraction of *BCL6* translocations (57% [27/47]) partnered to non-immunoglobulin loci (eg. *CIITA, RHOH, EIF4A2, ST6GAL1*; Table S7) compared to *BCL2* (1% [1/114]) and *MYC* (5% [2/38]) translocations (Figure 2A; Fisher P-value <0.001). These were more frequent in FL (88% [15/17] of *BCL6* translocations) as compared to DLBCL (39% [9/23] of *BCL6* translocations), presumably because the two immunoglobulin loci in FL are either translocated with the *BCL2* gene or functioning in immunoglobulin expression(21). We also employed off-target reads to detect DNA copy number alterations (CNAs) in a manner akin to low-pass whole genome sequencing, identified significant peaks of copy gain and losses using GISTIC2(18) (**Figures 1 and 2A**; Figure S4; Table S8-9), and defined the likely targets of these CNAs by integrative analysis of matched gene expression profiling (GEP) data from 290 tumors (**Figure 2B-C**, Figure S4, Table S10-11). This identified known CNA targets, including but not limited to deletion of *TNFAIP3* (6q24.2)(22), *ATM* (11q22.3)(23), *B2M* (15q15.5)(24) and *PTEN* (10q23.21)(25), and copy gain of *REL* and *BCL11A* (2p15), and *TCF4* (18q23)(26). In addition, we identified novel targets such as deletion of *IBTK* (6q14.1), *UBE3A* (11q22.1) and *FBXO25* (8p23.3), and copy gain of *ATF7* (12q13.13), *UCHL5* (1q31.3), and *KMT2A* (11q23.3). Importantly, the frequency of DNA copy number alterations in the target genes identified by NGS-based analysis significantly correlated with those derived from single nucleotide polymorphism (SNP) microarray-based measurements in independent cohorts of BL, DLBCL, FL and MCL tumors from previously published studies(6, 20, 26-30) (Figure S5), providing validation for the accuracy of this approach. The CNA peaks, defined as the smallest and most statistically significant region, included multiple genes that were significantly mutated (**Figure 2D**) as well as other genes for which we detected mutations at lower frequencies that were not significant by MutSig2CV (*POU2AF1, TP53BP1, FAS, PTEN*). Deletions of *ATM, B2M, BIRC3* and *TNFRSF14* significantly co-associated with mutations of these genes, suggesting that these are complementary mechanisms contributing to biallelic inactivation.

**Figure 2:**
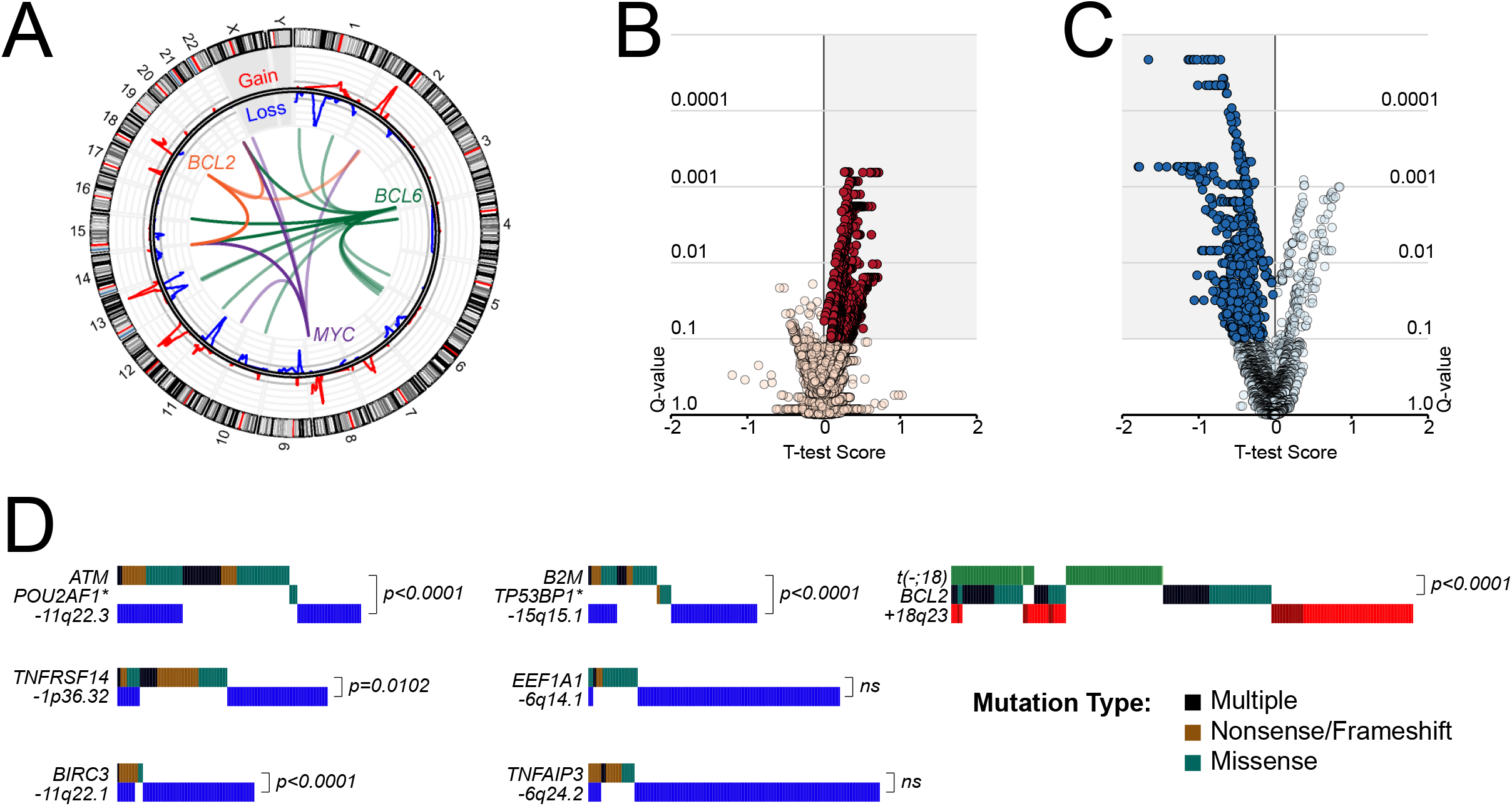
Structural alterations in B-NHL subtypes. **A)** A circos plot shows translocations of the MYC (purple), BCL2 (orange) and BCL6 (green) genes, and GISTIC tracks of DNA copy number gains (red) and losses (blue). **B-C)** Volcano plots of integrative analysis results showing the changes in gene expression of genes within peaks of DNA copy number gain (B) or loss (C). Positive T-test score indicate increased expression in tumors with a given CNA, and vice versa. Significant genes with the correct directionality are highlighted in the shaded areas. **D)** Oncoplots show the overlap of structural alterations and mutations that target the same genes. P-values are derived from a Fisher’s exact test (ns, not significant).

### Conserved functional hallmarks of B-NHL

To understand key hallmarks that are deregulated by genetic alterations, we performed hypergeometric enrichment analysis of genes targeted by recurrent mutations and DNA copy number alterations using DAVID(31) (Table S12). This revealed a significant enrichment of multiple overlapping gene sets that could be summarized into hallmark processes associated with epigenetics and transcription (**Figure 3A**), apoptosis and proliferation (**Figure 3B**), signaling (**Figure 3C**), and ubiquitination (**Figure 3D**). One or more genes within these hallmarks was altered in the majority (>50%) of tumors from each of the 4 major histologies included in this study. Genes annotated in epigenetic-associated gene sets were altered in 72%, 70%, 93% and 50% of BL, DLBCL, FL, and MCL, respectively, whereas genes annotated in transcription-associated gene sets were altered in 94%, 91%, 95% and 88% of BL, DLBCL, FL, and MCL, respectively. However, there is an extremely high degree of functional overlap between epigenetics and transcriptional regulation, as well as overlapping gene set annotations for many genes, leading us to consider these categories collectively as a single hallmark. Collectively, genes involved in epigenetics and transcription were mutated in 94% of BL, 92% of DLBCL, 96% of FL and 89% of MCL, and included those that encode proteins that catalyze post-translational modifications of histones (*KMT2D, CREBBP, EZH2, EP300, WHSC1, ASHL1L* and *KMT2A*), components of the BAF chromatin remodeling complex (*ARID1A, SMARCA4, BCL7A, BCL11A*), linker histones (*HIST1H1E, HIST1H1C, HIST1H1B*), and transcription factors (*BCL6, IRF4, IRF8, TCF3, TCF4, MYC, REL, PAX5, POU2AF2, FOXO1, CIITA*). Genes with a role in signaling included those involved in B-cell receptor (*CD79B, ID3, TCF3, TCF4, RFTN1*), NF_κ
_B (*TNFAIP3, CARD11, NFKBIE*), NOTCH (*NOTCH1, NOTCH2*), JAK/STAT (*SOCS1, STAT6*), PI3K/mTOR (*FOXO1, ATP6V1B2, APT6AP1*) and G-protein (GNA13, GNAI2) signaling. The *CD79A* and *BCL10* genes were also mutated at a lower frequency that was not significant by MutSig2CV (Figure S6A-B). Among these, the *RFTN1* gene (Figure S6C) is a novel recurrently mutated gene that was mutated in 7.4% of DLBCL and encodes a lipid raft protein that is critical for B-cell receptor signaling(32).

**Figure 3:**
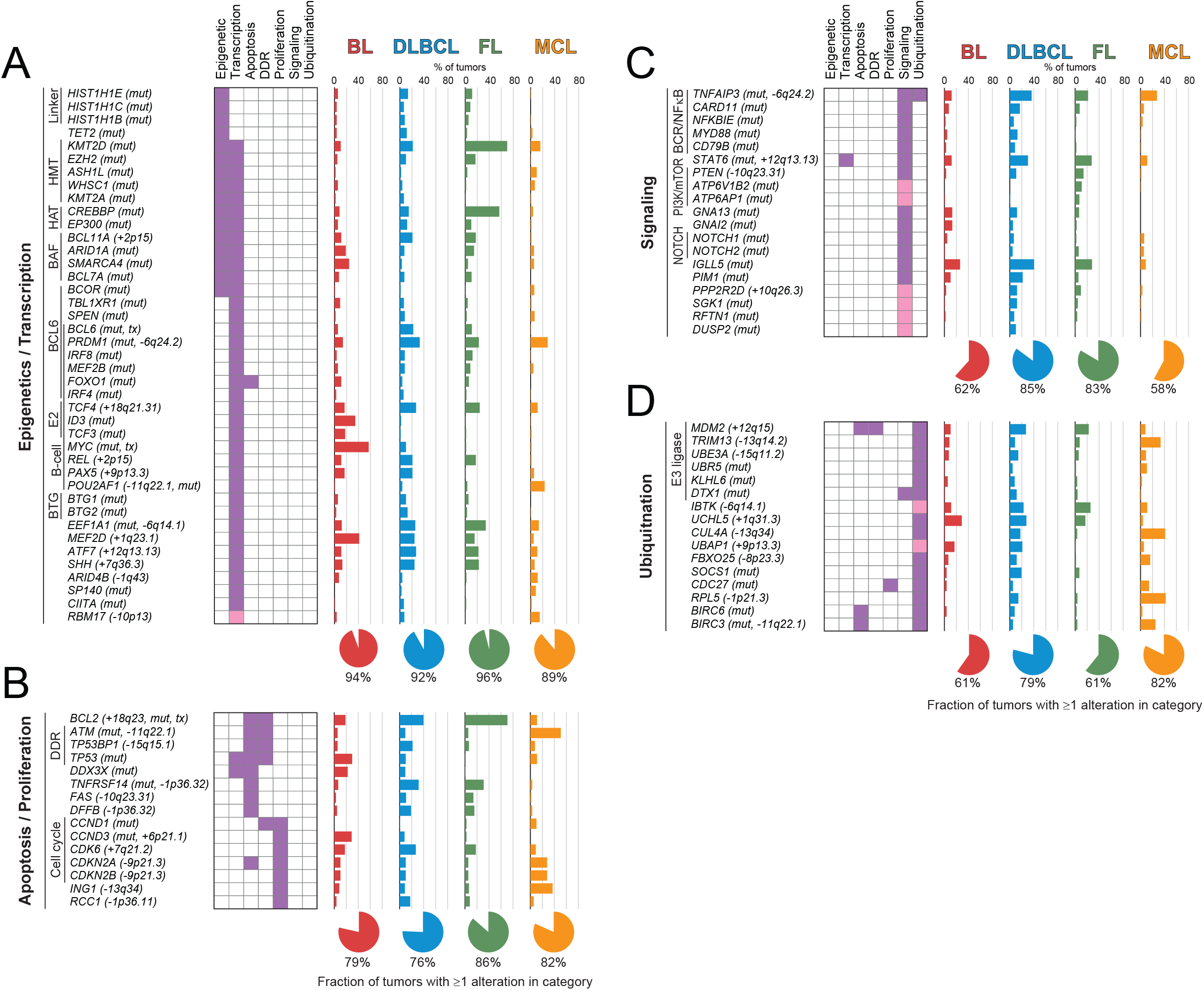
Functional enrichment of targets of somatic mutations and DNA copy number alterations. Genes targeted by somatic mutation and/or DNA copy number alteration were evaluated for enrichment in curated gene sets, and significant gene sets subsequently grouped according to overlapping gene set membership and functional similarity. In addition to genes assigned by DAVID (purple), some genes were manually curated into hallmark processes by literature review of their function (pink). Enriched gene sets could be summarized into 4 major hallmark processes, including (A) epigenetic and transcriptional control of gene expression, (B) regulation of apoptosis and proliferation, (C) regulation of signaling pathway activity, and (D) regulation of protein ubiquitination. The frequency of each genetic alteration is shown for each of the 4 major histologies included in this study, and the fraction of tumors in each histology bearing genetic alterations of one or more of the genes is summarized by a pie graph at the bottom for each hallmark. HMT, histone methyltransferase. HAT, histone acetyltransferase. DDR, DNA damage response. BCR, B-cell receptor.

Deregulation of the ubiquitin proteasome system is important in many cancers(33), but is not a well-defined hallmark of B-NHL. However, one or more genes with a role in regulating ubiquitination were genetically altered in 61% of BL, 79% of DLBCL, 61% of FL and 82% of MCL (**Figure 3D**). These included previously described genetic alterations such as amplification of *MDM2(34)*, deletion of *TNFAIP3(35)* and *CUL4A(36)* and *RPL5(36)*, and mutations of *KLHL6*(37), *DTX1(38), UBR5*(39), *SOCS1(40)*, and *BIRC3(6)*. In addition, we identified novel targets such as recurrent deletions of *IBTK*, a negative regulator of Bruton’s tyrosine kinase (BTK)(41), and somatic mutation of *CDC27* in 14% of MCL, which encodes an E3 ligase for CCND1(42). Therefore, common hallmark processes are targeted by genetic alterations in the majority of major B-NHL subtypes, including genes with a role in regulating protein ubiquitination.

### Subtype-specific patterns of genetic alterations

We formally tested the over- or under-representation of recurrent genetic alterations in each of the 4 subtypes with >100 tumors (BL, DLBCL, FL, MCL), compared to all other tumors in the study (**Figure 4**; Table S13). We observed some interesting patterns within hallmark characteristics that differ between subtypes. An illustrative example of this is the alternative avenues for BAF complex perturbation between different histologies (**Figure 5**). Specifically, mutations of the *SMARCA4* (aka. BRG1) component of the ATPase module were significantly enriched in BL (24%) compared to other subtypes (4%, Q-value<0.001), while mutations of the *BCL7A* component of the ATPase module were significantly enriched in FL (11%) compared to other subtypes (4%, Q-value=0.007). In contrast, mutations of *ARID1A* were frequent in both BL (19%) and FL (15%), and DNA copy number gains of *BCL11A* were frequent in both DLBCL (28%) and FL (22%). The BAF complex is therefore a target of recurrent genetic alterations, as previously suggested(43), but the manner in which this complex is perturbed varies between B-NHL subtypes (**Figure 5**). Similar disease-specific patterns were also observed for signaling genes; for example, *TCF3* and *ID3* have important functions in normal germinal center B-cells(44), but mutations of these genes are specifically enriched within BL and are rarely found in the other GCB-derived malignancies, DLBCL and FL. Similarly, the *ATP6AP1* and *ATP6V1B2* genes that function in mTOR signaling(45, 46) are specifically mutated in FL, and the *DUSP2* gene which inactivates ERK1/2(47) and STAT3(48) is specifically mutated in DLBCL. The disease-specific patterns of genetic alterations therefore reveal subtle but important differences in how each subtype of B-NHL perturbs hallmark features.

**Figure 4:**
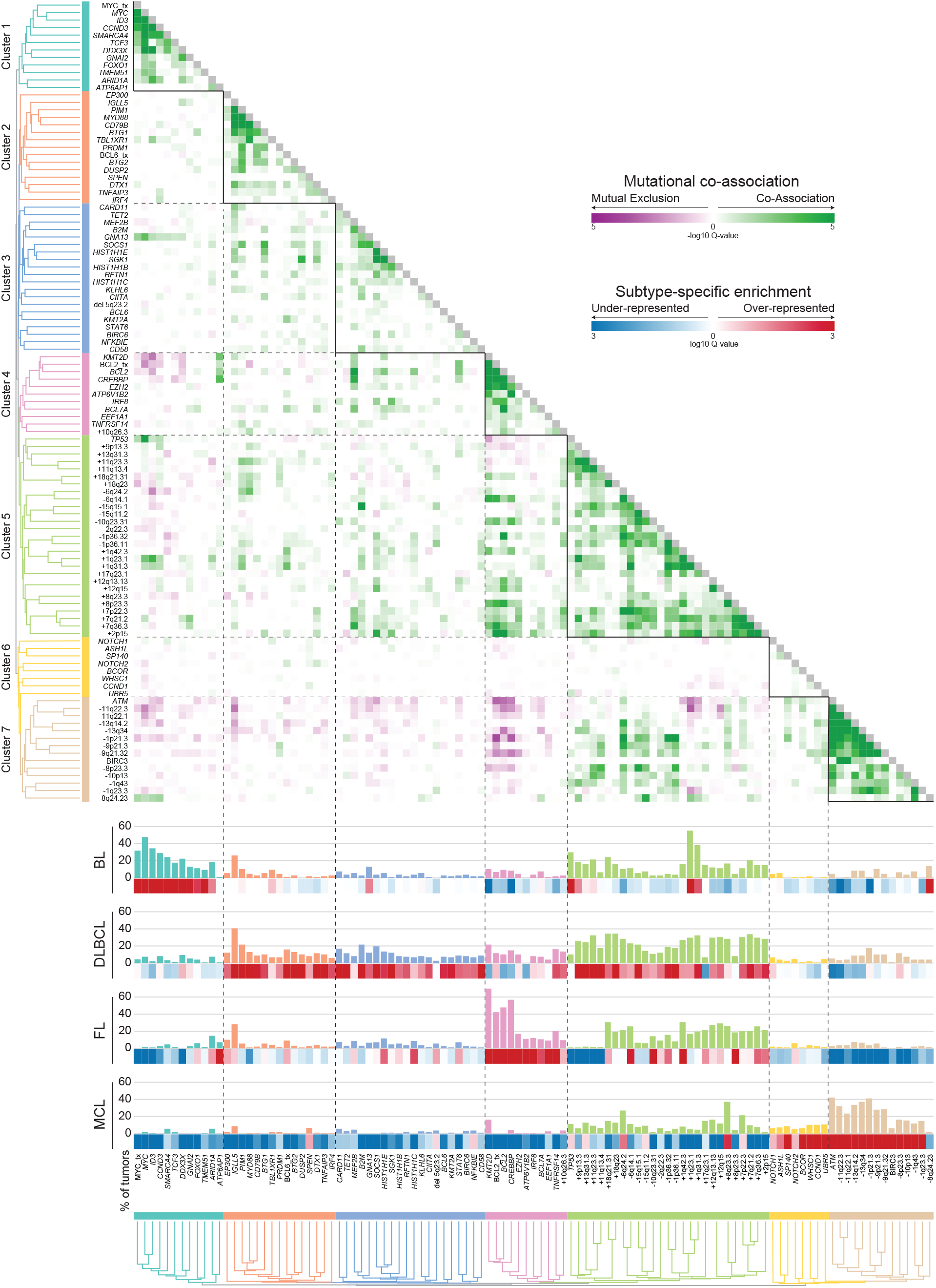
Subtype-specific clusters of co-occurring genetic alterations. The frequency (bar graph) and over/under-representation (blue to red scale) of mutations and structural alterations is shown on the left for BL, DLBCL, FL and MCL. The correlation matrix of co-associated (green) and mutually-exclusive (purple) relationships was clustered to identify 7 groups of co-occurring genetic alterations that were predominantly over-represented in a single B-NHL subtype.

**Figure 5:**
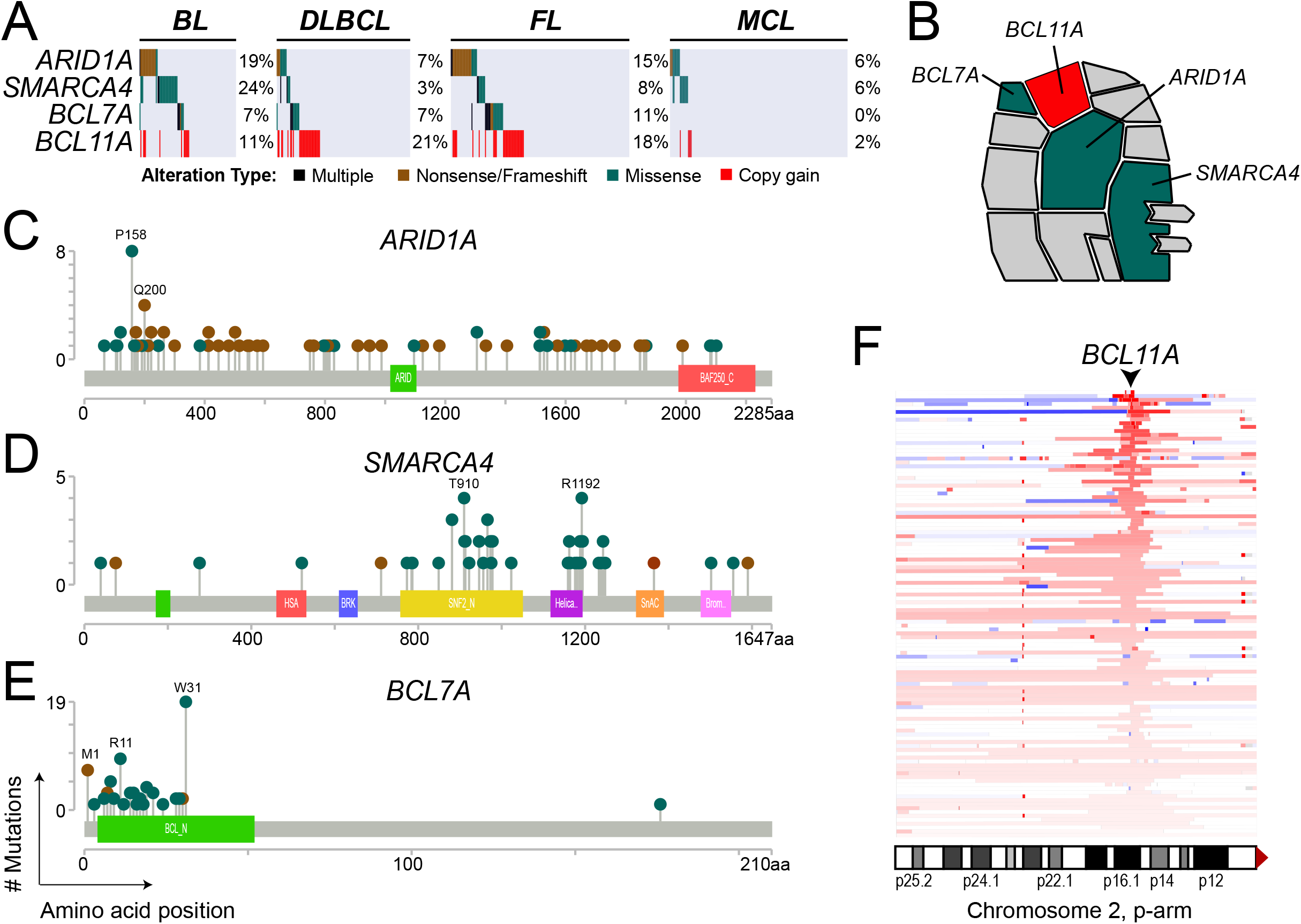
Subtype-specific patterns of BAF complex mutations. **A)** An oncoplot shows the frequency of genetic alterations in genes that encode components of the BAF complex. **B)** A schematic of the BAF complex shows recurrently mutated genes, *ARID1A, SMARCA4* and *BCL7A*, and the *BCL11A* gene that is targeted by 2p15 DNA copy number gains. **C-E)** Lollipop plots show the distribution of mutations in the BAF components *ARID1A* (C), *SMARCA4* (D), and *BCL7A* (E). **F)** A heatplot shows the location of chromosome 2p DNA copy number gains (red) ordered from highest DNA copy number (top) to lowest (bottom, copy number = 2.2). The *BCL11A* gene is in the peak focal copy gain.

### Clusters of co-associated genomic alterations in B-NHL subtypes

We next defined how each genetic alteration co-associated with or mutually-excluded other genetic alterations by pairwise assessments using false-discovery rate (FDR)-corrected Fisher’s tests (Table S14). A matrix of the transformed FDR Q-values (-logQ) was used for unsupervised hierarchical clustering to identify clusters of co-associated genetic alterations. Together with patterns of disease-specificity, unsupervised clustering revealed clear groupings of co-associated events for BL, DLBCL, FL and MCL (**Figure 4**). We identified a single cluster of significantly co-associated genetic alterations that was specifically enriched in BL (Cluster 1), including mutations and translocations of *MYC*, and mutations of *CCND3, SMARCA4, TCF3* and *ID3* that have been previously reported in BL(4). A single cluster was significantly enriched in MCL (Cluster 7), with a high frequency of *ATM* mutations and deletions, as well as other DNA copy number alterations. Other mutations that were not significantly co-associated were also enriched in MCL (Cluster 6), such as those in *WHSC1, NOTCH1, NOTCH2, BCOR* and *UBR5*, though statistical assessment of co-association may be hampered in this context by the low frequencies of mutations within these genes. A single cluster was also enriched in FL (Cluster 4), with a high prevalence of *KMT2D, BCL2, CREBBP, EZH2* and *TNFRSF14* mutations and *BCL2* translocations. The genes within cluster 4 also significantly overlapped with the previously reported C3, EZB and BCL2 clusters from prior whole exome sequencing studies of DLBCL (3, 49, 50) (Fisher test p-values = 0.0006, 0.0148 and 0.0173, respectively). Two clusters (Clusters 2 and 3) were enriched in DLBCL, with lower frequencies of mutations in a larger number of genes, in line with the genetic heterogeneity of this disease (3, 4). Cluster 2 includes co-associated genetic alterations that overlapped with the previously described C5, MCD, and MYD88 clusters (3, 4) (Fisher p-values = 0.0004, 0.0002 and 0.0007, respectively) including *CD79B, MYD88* and *TBL1XR1* mutations. Genes within cluster 3 significantly overlapped with those in the previously described C4 and SOCS1/SGK1 clusters (Fisher p-values = 0.0002 and 0.0074, respectively), including *SGK1, TET2, SOCS1* and histone H1 genes. We also identified a cluster consisting of TP53 mutations and multiple CNAs (Cluster 5) similar to the genetically complex C2/A53 subtype reported in DLBCL (3, 49), however the overlap of features within these clusters could not be formally assessed due to differing annotations. The CNAs captured in this cluster were variably represented across B-NHL subtypes, but were most frequent in DLBCL. B-NHL subtypes therefore harbor characteristic clusters of co-associated genetic alterations that likely cooperate in disease etiology.

### Combinations of genetic alterations define molecular subtypes of B-NHL

Our data have revealed statistical enrichment of individual genetic alterations in subtypes of B-NHL, and pairwise relationships between different genetic alterations that define clusters of subtype-specific events. To validate and expand upon these observations we leveraged gene expression microarray data from 284 tumors that underwent pathology review and were profiled as part of prior studies(10-12). We utilized BL, DHL, HGBL-NOS and DLBCL tumors to perform classification into molecular-defined BL (mBL) and non-mBL using a Bayesian classifier with previously described marker genes(51), and subclassified non-mBL into activated B-cell (ABC)-like and germinal center B-cell (GCB)-like subtypes as we have described previously(26) (Figure S7). We evaluated the frequency of cumulative (≥2) genetic alterations within each cluster among mBL, ABC-like DLBCL, GCB-like DLBCL, FL and MCL (**Figure 6**). This showed that Cluster 1 genetic alterations that were individually enriched in BL are cumulatively acquired in molecularly-defined BL (mBL), with 87% of tumors having ≥2 of these alterations compared to only 22% of GCB-like DLBCL. Similarly, Cluster 4 and Cluster 7 alterations were cumulatively in 77% and 72% of molecularly-annotated FL and MCL, respectively. Cluster 4 mutations were also cumulatively acquired in 51% of GCB-like DLBCL, likely capturing the C3/EZB/BCL2 subtype that has genetic similarities to FL(3, 4, 50). Furthermore, Cluster 2 and Cluster 4 alterations were cumulatively acquired in 58% of ABC-like DLBCL and 60% of GCB-like DLBCL, respectively, further supporting their respective overlap with the C5/MCD/MYD88 and C4/ST2 subtypes of DLBCL. CNAs within the Cluster 5 were cumulatively acquired at high but variable frequencies in all of the subtypes, but showed subtype-specific patterns within this cluster such as higher frequencies of 18q21 and 18q23 gains in ABC-like DLBCL, and higher frequencies of chromosome 7 gains in GCB-like DLBCL and FL. B-NHL tumors therefore cumulatively acquire co-associated sets of genetic alterations in a manner that is characteristically associated with histologically- and molecularly-defined subsets of disease.

**Figure 6:**
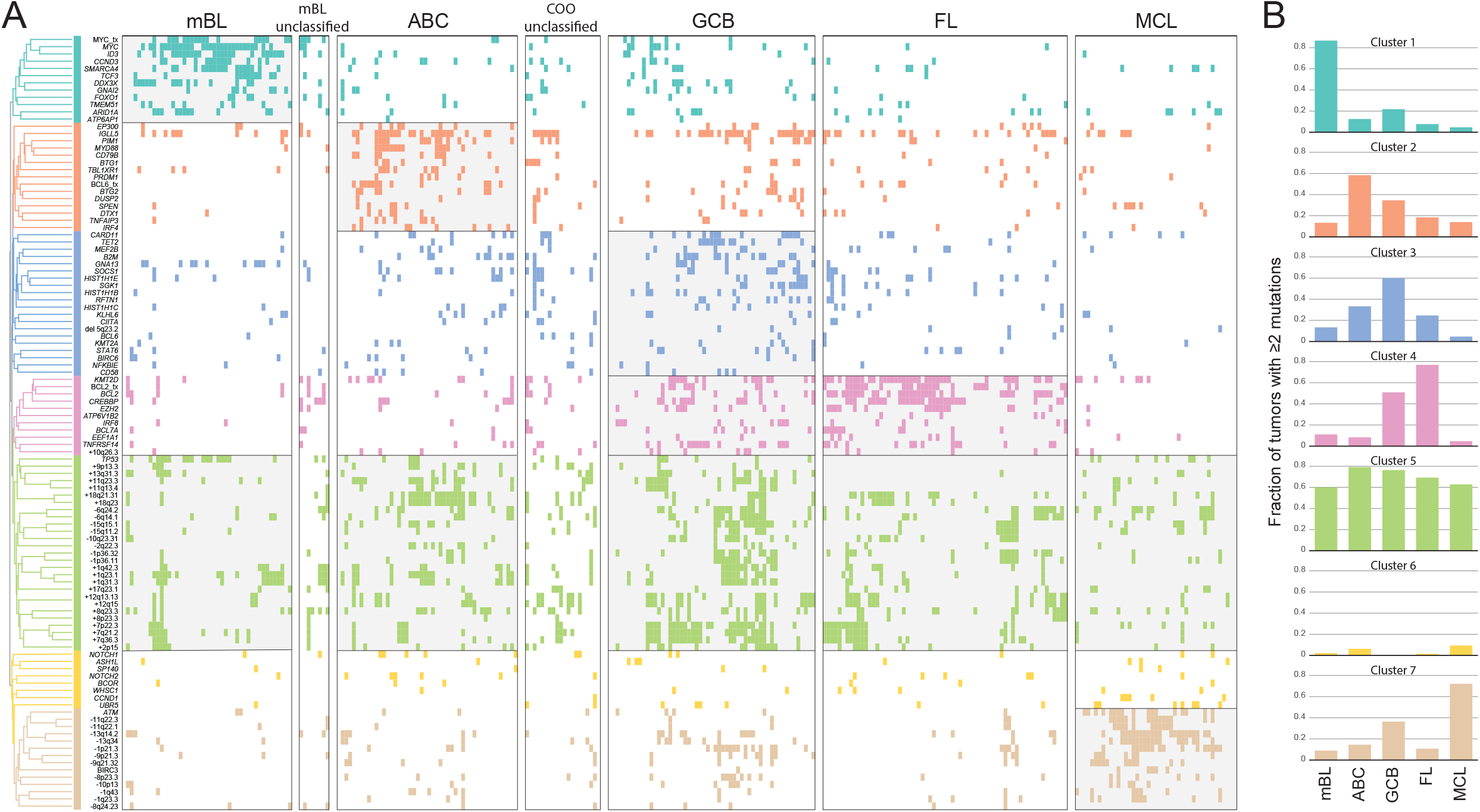
Cumulative acquisition of co-occurring genetic alterations. **A)** An oncoplot shows the presence or absence of genetic alterations according to their clusters of co-association in molecularly-defined Burkitt Lymphoma (mBL), activated B-cell (ABC)-like DLBCL, germinal center B-cell (GCB)-like DLBCL, FL and MCL with available gene expression microarray data. Shading shows histological or molecular subtypes with ≥50% of tumors bearing ≥2 genetic alterations within a given cluster. **B)** Bar plots shows the frequency of tumors with ≥2 genetic alterations from each cluster.

## DISCUSSION

By performing cross-sectional genomic profiling of a large cohort of tumors, we have developed a resource of genes and functional hallmarks that are recurrently targeted by genetic alterations in B-NHL, and shown that the cumulative acquisition of combinations of genetic alterations are characteristic of histological and molecular subtypes of disease. Some of the functional hallmarks that we identified have been previously appreciated, with a few exceptions. For example, the mutation of genes with roles in epigenetic and transcriptional control of gene expression are known to be a hallmark of FL(52) and we observed that 96% of FL tumors possessed mutations in one or more of the genes in this category. However, mutations within these genes were also observed in the majority of BL, DLBCL and MCL tumors, highlighting the conservation of this functional hallmark across B-NHL subtypes. There are subtype-specific patterns of chromatin modifying gene alterations, such as those that we highlighted for BAF complex mutations, but we suggest that the genetic deregulation of epigenetic and transcriptional control of gene expression should be considered a general hallmark of B-NHL. In addition, we suggest that the deregulation of the ubiquitin proteasome system is a hallmark of B-NHL that requires further investigation. Mutations in genes such as *KLHL6(37)* and *UBR5(39)* have been recently shown to play an important role in B-cell lymphoma, while the roles of other frequently mutated genes such as *DTX1* and *SOCS1* have not yet been functionally dissected. Furthermore, while the nature of AID-driven mutations in genes such as *DTX1* and *SOCS1* remain to be defined, other genes that are recurrently mutated by AID such as *BCL7A* (53) and linker histone genes (54) have been shown to play driving role in lymphomagenesis. Genetic deregulation of the ubiquitin proteasome system has the potential to influence the activity or abundance of a range of substrate proteins, and represents a current gap in our knowledge of B-NHL etiology.

The role of cooperative interactions between co-occurring genetic alterations is also an emerging field that requires further investigation. These interactions are not uncommon in cancer(55), and have been recently highlighted in DLBCL(3, 4), but our data show that they are pervasive and characteristic features of the B-NHL genetic landscape. Cooperation between co-associated genetic alterations identified in this study requires formal validation in cell line and/or animal models. However, there are many instances in which co-occurring genetic alterations that we observed have already been shown to cooperate in lymphomagenesis. In addition to the aforementioned example of *MYD88* and *CD79B* mutations, transgenic mouse models of *Ezh2* activating mutations or conditional deletion of *Crebbp* or *Kmt2d* have shown that these events are not alone sufficient for lymphomagenesis(56-61). We and others have observed a co-association between the mutation of these genes and *BCL2* translocations(14, 62), and the addition of a *Bcl2* transgene to these murine models indeed promoted lymphoma at a significantly higher rate than that observed with the *Bcl2* transgene alone(56-61). These genetic alterations are therefore significantly more lymphomagenic in combination than they are alone, which provides proof of principal that a cooperative relationship exists between these co-occurring genetic alterations. Future studies focusing on other co-occurring mutations, such as *MYC* translocation and *SMARCA4* mutation in BL, *CREBBP* and *KMT2D* mutation in FL, *TCF4* copy gain and *MYD88* mutation in DLBCL, and *ATM* mutation and *RPL5* deletion in MCL, should therefore be performed to further explore these concepts and define their underlying functional relationship. We suggest that combinations of genetic alterations are likely to more accurately recapitulate the biology of B-NHL than single gene models, and may reveal contextually different functional roles of genetic alterations depending on the co-occurring events.

The caveats of this study include the targeted nature of the LymphoSeq platform which may preclude consideration of a subset of important genes, the lack of germline DNA for the majority of samples that may lead to a small number of germ-line variants being falsely assigned as somatic, and the sample size for any given histological subtype being below that required to identify genes that are mutated at low frequency. Nonetheless, these data represent the first broad cross-sectional analysis of multiple histological and molecular subtypes of B-NHL using the same methodology and provide a framework of functional hallmarks and co-occurring genetic alterations that are enriched within these subtypes of B-NHL. These functional hallmarks are genetically perturbed in the majority of B-NHLs, but our cross-sectional approach enabled us to elucidate subtype-specific preferences for genetic alterations within each functional hallmark. Furthermore, the subtype-specific clusters of co-occurring genetic alterations likely represent cooperative interactions that underpin the biology of different subtypes of B-NHL. These combinations identify opportunities for moving from single-allele to multi-allele designs in cell line or animal models to better understand the molecular etiology of B-NHL subtypes. Together, these hallmarks and clusters of co-associated genetic alterations represent processes that are potentially drugable with targeted therapies (63-66), but that are likely influenced in a non-binary fashion by different combinations of genetic alterations. Deciphering the relationships between complex sets of genetic alterations and targetable dependencies will be a next step towards developing new rationally targeted therapeutic strategies in B-NHL.

## ACKNOWLEDGEMENTS

This research was supported by NCI R01CA201380 (MRG), the Nebraska Department of Health and Human Services (LB506 2016-17; MRG), and NCI cancer center support grants to the University of Texas MD Anderson Cancer Center (P30 CA016672) and the Fred & Pamela Buffet Cancer Center (P30 CA036727). HY is supported by a Fellow award from the Leukemia and Lymphoma Society. MRG is supported by a Scholar award from the Leukemia and Lymphoma Society and an Andrew Sabin Family Foundation Fellow award.

## AUTHOR CONTRIBUTIONS

MCJMA and ST performed experiments, analyzed data and wrote the manuscript. AB analyzed data. TH, HY, QD, DM, KH, NJ, JS, and SG performed experiments. AA, LS, MD, CC, JT, DP, KMV, MAL, ARS, BJC, RB, SN, LN, RED, JW, SP, MG, DS, KB, JI, SR, and AM provided samples and/or clinical data. MRG conceived and supervised the study, performed experiments, analyzed the data and wrote the manuscript. All authors reviewed and approved the manuscript.

## REFERENCES

1. Swerdlow SH, Campo E, Pileri SA, Harris NL, Stein H, Siebert R, et al. The 2016 revision of the World Health Organization classification of lymphoid neoplasms. Blood. 2016 May 19;127(20):2375–90.

2. Armitage JO, Gascoyne RD, Lunning MA, Cavalli F. Non-Hodgkin lymphoma. Lancet. 2017 Jul 15;390(10091):298–310.

3. Chapuy B, Stewart C, Dunford AJ, Kim J, Kamburov A, Redd RA, et al. Molecular subtypes of diffuse large B cell lymphoma are associated with distinct pathogenic mechanisms and outcomes. Nature medicine. 2018 Apr 30;24(5):679–90.

4. Schmitz R, Wright GW, Huang DW, Johnson CA, Phelan JD, Wang JQ, et al. Genetics and Pathogenesis of Diffuse Large B-Cell Lymphoma. The New England journal of medicine. 2018 Apr 12;378(15):1396–407.

5. Reddy A, Zhang J, Davis NS, Moffitt AB, Love CL, Waldrop A, et al. Genetic and Functional Drivers of Diffuse Large B Cell Lymphoma. Cell. 2017 Oct 05;171(2):481–94 e15.

6. Bea S, Valdes-Mas R, Navarro A, Salaverria I, Martin-Garcia D, Jares P, et al. Landscape of somatic mutations and clonal evolution in mantle cell lymphoma. Proceedings of the National Academy of Sciences of the United States of America. 2013 Nov 5;110(45):18250–5.

7. Zhang J, Jima D, Moffitt AB, Liu Q, Czader M, Hsi ED, et al. The genomic landscape of mantle cell lymphoma is related to the epigenetically determined chromatin state of normal B cells. Blood. 2014 May 08;123(19):2988–96.

8. Green MR, Alizadeh AA. Common progenitor cells in mature B-cell malignancies: implications for therapy. Current opinion in hematology. 2014 Jul;21(4):333–40.

9. Wang JQ, Jeelall YS, Humburg P, Batchelor EL, Kaya SM, Yoo HM, et al. Synergistic cooperation and crosstalk between MYD88L265P and mutations that dysregulate CD79B and surface IgM. The Journal of experimental medicine. 2017 Jul 12;214(9):2759–76.

10. Lenz G, Wright G, Dave SS, Xiao W, Powell J, Zhao H, et al. Stromal gene signatures in large-B-cell lymphomas. The New England journal of medicine. 2008 Nov 27;359(22):2313–23.

11. Dave SS, Fu K, Wright GW, Lam LT, Kluin P, Boerma EJ, et al. Molecular diagnosis of Burkitt’s lymphoma. The New England journal of medicine. 2006 Jun 8;354(23):2431–42.

12. Iqbal J, Shen Y, Liu Y, Fu K, Jaffe ES, Liu C, et al. Genome-wide miRNA profiling of mantle cell lymphoma reveals a distinct subgroup with poor prognosis. Blood. 2012 May 24;119(21):4939–48.

13. Bouska A, Bi C, Lone W, Zhang W, Kedwaii A, Heavican T, et al. Adult High Grade B-cell Lymphoma with Burkitt Lymphoma Signature: Genomic features and Potential Therapeutic Targets. Blood. 2017 Aug 11.

14. Green MR, Kihira S, Liu CL, Nair RV, Salari R, Gentles AJ, et al. Mutations in early follicular lymphoma progenitors are associated with suppressed antigen presentation. Proceedings of the National Academy of Sciences of the United States of America. 2015 Mar 10;112(10):E1116–25.

15. Lek M, Karczewski KJ, Minikel EV, Samocha KE, Banks E, Fennell T, et al. Analysis of protein-coding genetic variation in 60,706 humans. Nature. 2016 Aug 18;536(7616):285–91.

16. Lawrence MS, Stojanov P, Polak P, Kryukov GV, Cibulskis K, Sivachenko A, et al. Mutational heterogeneity in cancer and the search for new cancer-associated genes. Nature. 2013 Jul 11;499(7457):214–8.

17. Kuilman T, Velds A, Kemper K, Ranzani M, Bombardelli L, Hoogstraat M, et al. CopywriteR: DNA copy number detection from off-target sequence data. Genome biology. 2015;16:49.

18. Mermel CH, Schumacher SE, Hill B, Meyerson ML, Beroukhim R, Getz G. GISTIC2.0 facilitates sensitive and confident localization of the targets of focal somatic copy-number alteration in human cancers. Genome biology. 2011;12(4):R41.

19. Newman AM, Bratman SV, Stehr H, Lee LJ, Liu CL, Diehn M, et al. FACTERA: a practical method for the discovery of genomic rearrangements at breakpoint resolution. Bioinformatics. 2014 Dec 1;30(23):3390–3.

20. Bouska A, Bi C, Lone W, Zhang W, Kedwaii A, Heavican T, et al. Adult high-grade B-cell lymphoma with Burkitt lymphoma signature: genomic features and potential therapeutic targets. Blood. 2017 Oct 19;130(16):1819–31.

21. Akasaka T, Lossos IS, Levy R. BCL6 gene translocation in follicular lymphoma: a harbinger of eventual transformation to diffuse aggressive lymphoma. Blood. 2003 Aug 15;102(4):1443–8.

22. Kato M, Sanada M, Kato I, Sato Y, Takita J, Takeuchi K, et al. Frequent inactivation of A20 in B-cell lymphomas. Nature. 2009 Jun 04;459(7247):712–6.

23. Greiner TC, Dasgupta C, Ho VV, Weisenburger DD, Smith LM, Lynch JC, et al. Mutation and genomic deletion status of ataxia telangiectasia mutated (ATM) and p53 confer specific gene expression profiles in mantle cell lymphoma. Proceedings of the National Academy of Sciences of the United States of America. 2006 Feb 14;103(7):2352–7.

24. Challa-Malladi M, Lieu YK, Califano O, Holmes AB, Bhagat G, Murty VV, et al. Combined genetic inactivation of beta2-Microglobulin and CD58 reveals frequent escape from immune recognition in diffuse large B cell lymphoma. Cancer cell. 2011 Dec 13;20(6):728–40.

25. Pfeifer M, Grau M, Lenze D, Wenzel SS, Wolf A, Wollert-Wulf B, et al. PTEN loss defines a PI3K/AKT pathway-dependent germinal center subtype of diffuse large B-cell lymphoma. Proceedings of the National Academy of Sciences of the United States of America. 2013 Jul 23;110(30):12420–5.

26. Jain N, Hartert K, Tadros S, Fiskus W, Havranek O, Ma M, et al. Targetable genetic alterations of TCF4 (E2-2) drive immunoglobulin expression in the activated B-cell subtype of diffuse large B-cell lymphoma. Sci Transl Med. 2019;11(497):eeav5599.

27. Kim D, Fiske BP, Birsoy K, Freinkman E, Kami K, Possemato RL, et al. SHMT2 drives glioma cell survival in ischaemia but imposes a dependence on glycine clearance. Nature. 2015 Apr 16;520(7547):363–7.

28. Rosenwald A, Wright G, Wiestner A, Chan WC, Connors JM, Campo E, et al. The proliferation gene expression signature is a quantitative integrator of oncogenic events that predicts survival in mantle cell lymphoma. Cancer cell. 2003 Feb;3(2):185–97.

29. Salaverria I, Royo C, Carvajal-Cuenca A, Clot G, Navarro A, Valera A, et al. CCND2 rearrangements are the most frequent genetic events in cyclin D1(-) mantle cell lymphoma. Blood. 2013 Feb 21;121(8):1394–402.

30. Green MR, Vicente-Duenas C, Romero-Camarero I, Long Liu C, Dai B, Gonzalez-Herrero I, et al. Transient expression of Bcl6 is sufficient for oncogenic function and induction of mature B-cell lymphoma. Nature communications. 2014;5:3904.

31. Huang da W, Sherman BT, Lempicki RA. Systematic and integrative analysis of large gene lists using DAVID bioinformatics resources. Nature protocols. 2009;4(1):44–57.

32. Saeki K, Miura Y, Aki D, Kurosaki T, Yoshimura A. The B cell-specific major raft protein, Raftlin, is necessary for the integrity of lipid raft and BCR signal transduction. EMBO J. 2003 Jun 16;22(12):3015–26.

33. Senft D, Qi J, Ronai ZA. Ubiquitin ligases in oncogenic transformation and cancer therapy. Nature reviews Cancer. 2018 Feb;18(2):69–88.

34. Monti S, Chapuy B, Takeyama K, Rodig SJ, Hao Y, Yeda KT, et al. Integrative analysis reveals an outcome-associated and targetable pattern of p53 and cell cycle deregulation in diffuse large B cell lymphoma. Cancer cell. 2012 Sep 11;22(3):359–72.

35. Honma K, Tsuzuki S, Nakagawa M, Tagawa H, Nakamura S, Morishima Y, et al. TNFAIP3/A20 functions as a novel tumor suppressor gene in several subtypes of non-Hodgkin lymphomas. Blood. 2009 Sep 17;114(12):2467–75.

36. Hartmann EM, Campo E, Wright G, Lenz G, Salaverria I, Jares P, et al. Pathway discovery in mantle cell lymphoma by integrated analysis of high-resolution gene expression and copy number profiling. Blood. 2010 Aug 12;116(6):953–61.

37. Choi J, Lee K, Ingvarsdottir K, Bonasio R, Saraf A, Florens L, et al. Loss of KLHL6 promotes diffuse large B-cell lymphoma growth and survival by stabilizing the mRNA decay factor roquin2. Nat Cell Biol. 2018 May;20(5):586–96.

38. Meriranta L, Pasanen A, Louhimo R, Cervera A, Alkodsi A, Autio M, et al. Deltex-1 mutations predict poor survival in diffuse large B-cell lymphoma. Haematologica. 2017 May;102(5):e195–e8.

39. Swenson SA, Gilbreath TJ, Vahle H, Hynes-Smith RW, Graham JH, Law HC, et al. UBR5 HECT domain mutations identified in mantle cell lymphoma control maturation of B cells. Blood. 2020 Jul 16;136(3):299–312.

40. Mottok A, Renne C, Seifert M, Oppermann E, Bechstein W, Hansmann ML, et al. Inactivating SOCS1 mutations are caused by aberrant somatic hypermutation and restricted to a subset of B-cell lymphoma entities. Blood. 2009 Nov 12;114(20):4503–6.

41. Liu W, Quinto I, Chen X, Palmieri C, Rabin RL, Schwartz OM, et al. Direct inhibition of Bruton’s tyrosine kinase by IBtk, a Btk-binding protein. Nature immunology. 2001 Oct;2(10):939–46.

42. Pawar SA, Sarkar TR, Balamurugan K, Sharan S, Wang J, Zhang Y, et al. C/EBP{delta} targets cyclin D1 for proteasome-mediated degradation via induction of CDC27/APC3 expression. Proceedings of the National Academy of Sciences of the United States of America. 2010 May 18;107(20):9210–5.

43. Krysiak K, Gomez F, White BS, Matlock M, Miller CA, Trani L, et al. Recurrent somatic mutations affecting B-cell receptor signaling pathway genes in follicular lymphoma. Blood. 2017 Jan 26;129(4):473–83.

44. Gloury R, Zotos D, Zuidscherwoude M, Masson F, Liao Y, Hasbold J, et al. Dynamic changes in Id3 and E-protein activity orchestrate germinal center and plasma cell development. The Journal of experimental medicine. 2016 May 30;213(6):1095–111.

45. Okosun J, Wolfson RL, Wang J, Araf S, Wilkins L, Castellano BM, et al. Recurrent mTORC1-activating RRAGC mutations in follicular lymphoma. Nature genetics. 2016 Feb;48(2):183–8.

46. Wang F, Gatica D, Ying ZX, Peterson LF, Kim P, Bernard D, et al. Follicular lymphoma-associated mutations in vacuolar ATPase ATP6V1B2 activate autophagic flux and mTOR. The Journal of clinical investigation. 2019 Mar 4;130:1626–40.

47. Hu J, Li L, Chen H, Zhang G, Liu H, Kong R, et al. MiR-361-3p regulates ERK1/2-induced EMT via DUSP2 mRNA degradation in pancreatic ductal adenocarcinoma. Cell Death Dis. 2018 Jul 24;9(8):807.

48. Lu D, Liu L, Ji X, Gao Y, Chen X, Liu Y, et al. The phosphatase DUSP2 controls the activity of the transcription activator STAT3 and regulates TH17 differentiation. Nature immunology. 2015 Dec;16(12):1263–73.

49. Wright GW, Huang DW, Phelan JD, Coulibaly ZA, Roulland S, Young RM, et al. A Probabilistic Classification Tool for Genetic Subtypes of Diffuse Large B Cell Lymphoma with Therapeutic Implications. Cancer cell. 2020 Apr 13;37(4):551–68 e14.

50. Lacy SE, Barrans SL, Beer PA, Painter D, Smith AG, Roman E, et al. Targeted sequencing in DLBCL, molecular subtypes, and outcomes: a Haematological Malignancy Research Network report. Blood. 2020 May 14;135(20):1759–71.

51. Hummel M, Bentink S, Berger H, Klapper W, Wessendorf S, Barth TF, et al. A biologic definition of Burkitt’s lymphoma from transcriptional and genomic profiling. The New England journal of medicine. 2006 Jun 8;354(23):2419–30.

52. Green MR. Chromatin modifying gene mutations in follicular lymphoma. Blood. 2018 Feb 8;131(6):595–604.

53. Balinas-Gavira C, Rodriguez MI, Andrades A, Cuadros M, Alvarez-Perez JC, Alvarez-Prado AF, et al. Frequent mutations in the amino-terminal domain of BCL7A impair its tumor suppressor role in DLBCL. Leukemia. 2020 Oct;34(10):2722–35.

54. Yusufova N, Kloetgen A, Teater M, Osunsade A, Camarillo JM, Chin CR, et al. Histone H1 loss drives lymphoma by disrupting 3D chromatin architecture. Nature. 2020 Dec 9.

55. Ashworth A, Lord CJ, Reis-Filho JS. Genetic interactions in cancer progression and treatment. Cell. 2011 Apr 1;145(1):30–8.

56. Beguelin W, Popovic R, Teater M, Jiang Y, Bunting KL, Rosen M, et al. EZH2 is required for germinal center formation and somatic EZH2 mutations promote lymphoid transformation. Cancer cell. 2013 May 13;23(5):677–92.

57. Garcia-Ramirez I, Tadros S, Gonzalez-Herrero I, Martin-Lorenzo A, Rodriguez-Hernandez G, Moore D, et al. Crebbp loss cooperates with Bcl2 over-expression to promote lymphoma in mice. Blood. 2017 Mar 13;129(19):2645–56.

58. Zhang J, Vlasevska S, Wells VA, Nataraj S, Holmes AB, Duval R, et al. The Crebbp Acetyltransferase is a Haploinsufficient Tumor Suppressor in B Cell Lymphoma. Cancer discovery. 2017 Jan 09;7(3):322–37.

59. Jiang Y, Ortega-Molina A, Geng H, Ying HY, Hatzi K, Parsa S, et al. CREBBP Inactivation Promotes the Development of HDAC3-Dependent Lymphomas. Cancer discovery. 2017 Jan;7(1):38–53.

60. Zhang J, Dominguez-Sola D, Hussein S, Lee JE, Holmes AB, Bansal M, et al. Disruption of KMT2D perturbs germinal center B cell development and promotes lymphomagenesis. Nature medicine. 2015 Oct;21(10):1190–8.

61. Ortega-Molina A, Boss IW, Canela A, Pan H, Jiang Y, Zhao C, et al. The histone lysine methyltransferase KMT2D sustains a gene expression program that represses B cell lymphoma development. Nature medicine. 2015 Oct;21(10):1199–208.

62. Morin RD, Mendez-Lago M, Mungall AJ, Goya R, Mungall KL, Corbett RD, et al. Frequent mutation of histone-modifying genes in non-Hodgkin lymphoma. Nature. 2011 Aug 18;476(7360):298–303.

63. Sermer D, Pasqualucci L, Wendel HG, Melnick A, Younes A. Emerging epigenetic-modulating therapies in lymphoma. Nature reviews Clinical oncology. 2019 Aug;16(8):494–507.

64. Shen M, Schmitt S, Buac D, Dou QP. Targeting the ubiquitin-proteasome system for cancer therapy. Expert Opin Ther Targets. 2013 Sep;17(9):1091–108.

65. Merino D, Kelly GL, Lessene G, Wei AH, Roberts AW, Strasser A. BH3-Mimetic Drugs: Blazing the Trail for New Cancer Medicines. Cancer cell. 2018 Dec 10;34(6):879–91.

66. Roschewski M, Staudt LM, Wilson WH. Diffuse large B-cell lymphoma-treatment approaches in the molecular era. Nature reviews Clinical oncology. 2014 Jan;11(1):12–23.

