## Supplementary Methods for "Subtype-specific and co-occurring genetic alterations in B-cell non-Hodgkin lymphoma"

Man Chun John Ma^¥^, Saber Tadros^¥^, Alyssa Bouska, Tayla Heavican, Haopeng Yang, Qing Deng, Dalia Moore, Ariz Akhter, Keenan Hartert, Neeraj Jain, Jordan Showell, Sreejoyee Ghosh, Lesley Street, Marta Davidson, Christopher Carey, Joshua Tobin, Deepak Perumal, Julie M. Vose, Matthew A. Lunning, Aliyah R. Sohani, Benjamin J. Chen, Shannon Buckley, Loretta J. Nastoupil, R. Eric Davis, Jason R. Westin, Nathan H. Fowler, Samir Parekh, Maher Gandhi, Sattva Neelapu, Douglas Stewart, Kapil Bhalla, Javeed Iqbal, Timothy Greiner, Scott J. Rodig, Adnan Mansoor, Michael R. Green*

**CONTENTS**

Supplementary Methods

Figure S1: Overview of approach

Figure S2: Validation of CopyWriteR DNA copy number calls.

Figure S3: Spectrum of mutational burden and copy number aberrant genome

Figure S4: GISTIC and integrative analysis

Figure S5: Comparison of DNA copy number alteration frequencies between NGS- and SNP microarray-based copy number analysis

Figure S6: Lollipop plots for select genes

Figure S7: Molecular classification of UNMC tumors

Supplementary References

Supplementary Tables

Table S1: Sample information

Table S2: LymphoSeq target genes

Table S3: Coding SNVs and InDels

Table S4: MutSig2CV results

Table S5: Gene-level frequency of mutations in AID motifs

Table S6: Mutational signatures by disease subtype

Table S7: Translocations of MYC, BCL2 and BCL6

Table S8: Genes in GISTIC2 copy gain peaks

Table S9: Genes in GISTIC2 copy loss peaks

Table S10: Integrative analysis of copy gains

Table S11: Integrative analysis of copy losses

Table S12: DAVID hypergeometric enrichment analysis

Table S13: Fisher tests of disease association

Table S14: Fisher tests of mutual exclusivity and co-occurrence

Table S15: Recurrent false positive genes

**SUPPLEMENTARY METHODS**

**Target Selection for LymphoSeq**

A catalogue of somatic mutations in 670 B-cell malignancies was compiled from previous high-throughput sequencing studies with sufficient quality and variant reporting, and with germline controls, that were available during the design phase in 2014. These included 53 Mantle cell lymphoma (MCL)^1-3^, 82 Burkitt’s lymphoma (BL)^4,5^, 155 Diffuse large B-cell lymphoma (DLBCL)^6-9^, 65 Follicular lymphoma (FL)^6,10-13^, 214 Chronic lymphocytic leukemia (CLL)^14-17^, 36 Marginal zone lymphoma^18,19^, 38 Multiple myeloma^20^ (MM), 10 Waldenstrom’s macroglobulinemia (WM)^21^, 9 Primary mediastinal large B-cell lymphoma^22^ (PMBCL), 8 Primary central nervous system lymphoma^23^ (PCNSL). Genes that were mutated in >2% of any individual subtype (with WM, PMBCL and PCNSL being grouped together as rare lymphomas), and that were not previously identified as genes with recurrent false-positive mutation calls^10^ (MUC4, RPTN, PRH1, DSPP, C9ORF96, NBPF1, MUC6, MAML2, MAN1B1, C9ORF84, PDE4DIP, PTPRN2, GPC1, TBP, PKP2, KIAA1683, TTC7B, OVOS2), were selected for targeting by hybrid capture. This resulted in the coding and untranslated regions of 380 genes (Table S2) totaling 3,232,925 bp.

**Detailed Next Generation Sequencing Methods**

Library Preparation. Next generation sequencing libraries were prepared from 100 – 1000ng of genomic DNA (gDNA). High molecular weight gDNA was sheared by sonication in a Covaris S220 or M220 instrument (Covaris Inc.) to obtain an average molecular weight of 150-200bp. Sheared DNA was cleaned up with Ampure Beads (Beckman Coulter), and libraries prepared using Hyper Prep kits (KAPA Biosystems), according to the manufacturer’s protocol except for an overnight ligation of adapters at 16˚C. A maximum of 8 cycles of PCR was performed during library preparation (mean = 6 cycles). TruSeq adapters (Bioo Scientific) were utilized at the recommended ratio to input DNA. Library quality was assessed using TapeStation High-Sensitivity DNA 1000 (Agilent) and quantified by Qubit (Life Technologies).

Hybrid capture and sequencing. Libraries were 12-plexed in equal quantities and a 1µg of pooled libraries were enriched by hybrid capture using a Nimblegen SeqCap EZ Custom reagent (Roche), according to the manufacturer’s protocol. Each capture pool was sequenced on a single lane of a HiSeq 2500 in high output mode using 2 x 100bp at the Hudson Alpha Institute for Biotechnology. A total of 2.06 Tera-base-pairs of sequencing data was produced. The average on-target rate and sequencing depth of the 685 samples presented in this study were 81.3% (range, 71.7% – 92.5%) and 624X (range, 101X – 1785X), respectively, determined as per below.

Variant calling. Raw FASTQ files were assessed for quality using FASTQC. Samples with high quality metrics were run through our in-house pipeline. FASTQ files were (i) aligned to the human genome (hg19) using BWA-Mem^24^, (ii) deduplicated using Picard MarkDuplicates, (iii) realigned around InDels using GATK^25^, and (iv) recalibrated by base score using GATK^25^. On-target rate and coverage over the targeted region were calculated by Picard CalculateHSMetrics. Only samples with >100X average coverage were utilized. If additional sequencing was required to obtain sufficient coverage, bam files from the same sample were merged using BamTools Mappings Merger following alignment. Variants were called by GATK Unified Genotyper and VarScan2. Only variants called by both tools were retained, which we have previously shown to provide a sensitivity of 96.7% and specificity of 92.9%^11^. All variants were annotated using SeattleSeq^26^.

AID mutations. Mutations were defined as products of AID activity if the wild-type allele was a cytosine within the context of a WRCY motif, as previously described^11^. These data are reported in Table S5 as the total number of mutations in the coding sequence of each gene that fit this criteria.

Filtering of repetitive regions and potential germline variants. To avoid mapping artifacts in repetitive regions, all variants within RepeatMasker or tandemRepeat annotated regions were filtered from the dataset. All genes selected for capture had prior evidence of somatic mutation from whole exome or whole genome sequencing studies (see ‘Target Selection for Hybrid Capture’ section). In addition, all variants in dbSNP build 32 were removed, and all mutation hotspots (defined below) were manually searched in the ExAC browser^27^ containing whole exome sequencing of 60,706 individuals. Genes with mutation hotspots present in ≥1 healthy individual were filtered from the dataset to remove the chance of these variants corresponding to a germline polymorphism. Mutation frequencies of significantly mutated genes (MutSigCV, below) were compared between fresh/frozen and FFPE tumors for BL and MCL using a Fisher exact test. No significant differences were detected, therefore filtering was not performed using this criteria.

MutSig2CV. MAF files containing filtered variants were analyzed by MutSig2CV^28^. This analysis was performed for the data set as a whole, and for the four major subtypes individually (DLBCL, BL, FL and MCL). Genes that reached a significance of Q<0.25 in any one of these analyses were included, as well as genes that did not reach this threshold but have a well-defined role in lymphoma and are targets of somatic hypermutation (eg. *BCL2* and *BCL6*; Table S5).

DNA copy number and GISTIC2 analysis. Genome-wide DNA copy number profiles were determined from mapped, realigned, sorted, deduplicated bam files using CopyWriteR^29^. For the generation of CopywriteR library folder we utilized 100kb bins in hg19 assembly with contigs with a prefix of 'chr' [preCopywriteR (bin.size=100000, ref.genome='hg19', prefix='chr')]. For the identification of CNAs we used “input vs. none” [CopywriteR(sample.control, destination.folder, reference.folder, bp.param, capture.regions.file)], where sample.control is generated by data.frame(sample=[input_bam], control=[input_bam]), reference.folder is the folder generated by preCopywriteR above, bp.param involves parallelism, and capture.regions.file is the BED file containing the LymphoSeq panel's capture regions. Segmented data were analyzed using GISTIC2^30^ in the GenePattern^31^ environment, with copy number thresholds of 0.2 and marker number of 100. Peaks were considered to be significant if they had a residual Q-value < 0.1, to account for the significance in neighboring peaks. The gene-level data from GISTIC2 analysis have been uploaded to cBioPortal. The genes within each peak were used for integrative analysis with matched gene expression profiling data (below) to define the likely drivers of each alteration. For validation, DNA copy number alterations of genes shown in Figure 3 were evaluated in previously reported SNP microarray data from 85 BL, 694 DLBCL, 404 FL and 206 MCL. For BL^32^, DLBCL^33,34^ and FL^35^, the combined analysis of these datasets have been previously reported. For MCL, high-resolution (>200,000 markers) Affymetrix SNP microarray or Agilent CGH array DNA number data were downloaded for 206 tumors from the gene expression omnibus (GSE12906, GSE18820, GSE42854, GSE46969) and probe-level data segmented using the cbs tool in GenePattern^31^. Gene level DNA copy number was calculated from segmented data using GISTIC2, and reported as copy loss or gain if absolute DNA copy number was <1.8 or >2.2, respectively, in line with the thresholds used for GISTIC analysis of the NGS cohort. Heatmaps were generated using the GENE-E tool.

Identification of B-NHL hallmarks. A gene list was created that included all genes that were significantly mutated, as determined by MutSig2CV, and/or targeted by DNA copy number gain or loss, as determined by GISTIC2 and integrative analysis. This gene list was interrogated using the ‘Functional Annotation Clustering’ function of DAVID^36^. This identified 41 clusters of significantly enriched pathways, which were grouped together according to functional relatedness.

Assessment of co-association and clustering. Genetic alterations that were present in ≥5% of any major histologic subtype (BL, DLBCL, FL, MCL) were assessed for mutual co-association or exclusion using a Fisher exact test corrected for multiple hypothesis testing by the Benjamini-Hochberg method (Table S14). Co-association q-values were transformed by log10 and directionality (co-association assigned as positive values, exclusion as negative values) and then a matrix of transformed values was clustered in GENE-E using Pearson’s correlation coefficient with complete linkage. Genes within the clusters reported in this study were compared to those from other studies (Lacy, Chapuy, Wright) using a Fisher exact test. DNA copy number alterations were omitted for this comparison due to absence (Lacy) or differences in the nomenclature used (Wright).

**Gene Expression Analysis**

Affymetrix U133 plus 2.0 gene expression microarray data were available for 284 tumors from the University of Nebraska Medical Center (Table S1)^37-39^. Raw cel files were RMA normalized using the ExpressionFileCreator module of GenePattern^31^, and quality checked for batch effects by unsupervised clustering of the 3,000 most variably expressed genes across the dataset.

Integrative analysis of DNA copy number alterations. Integrative analysis was performed using tumor subsets with ≥3 tumors with a given copy number alterations. If tumor subtypes did not harbor the copy number alteration of interest, they were excluded so as not to confound the analysis with subtype-specific gene expression differences. For each DNA copy number alteration, the expression of genes within the peak was compared between diploid and gain/loss tumors using a Student’s T-test corrected for multiple hypothesis testing with a Benjamini-Hochberg correction. Genes with a directionality of change matching the DNA copy number alteration type (i.e. increased expression with DNA copy gain, decreased expression with DNA copy loss) and a Q-value < 0.25 were considered significant.

Molecular Burkitt lymphoma (mBL) classification. For mBL classification, we built a Bayesian classifier using previously published dataset^40^. The model was created on the discovery set from the original paper and evaluated on the test set. Classifications based upon our model had minor differences to the originally reported classification, primarily in reducing the number of unclassified tumors, and improved upon the significance in overall survival difference between mBL and non-mBL subgroups (Figure S6). We therefore applied this model to the UNMC dataset, including DLBCL, BL and HGBL-NOS tumors (n=159).

Cell of origin subtyping. For COO subtyping, we utilized a previously described 140 gene Bayesian classifier^41^. This classification was applied to non-mBL tumors from the molecular Burkitt classifier, and overall survival assessed in the subgroups as a sanity check (Figure S6).

**Data Sharing**. All gene expression data has been consolidated under a single Gene Expression Omnibus Accession (GSE132929). Mutation, DNA copy number alteration, translocation and gene expression data have been uploaded to cBioPortal for easy access (<https://www.cbioportal.org/study/summary?id=mbn_mdacc_2013>). Raw next generation sequencing data will be made available to investigators at academic or not-for-profit research institutions upon reasonable request.

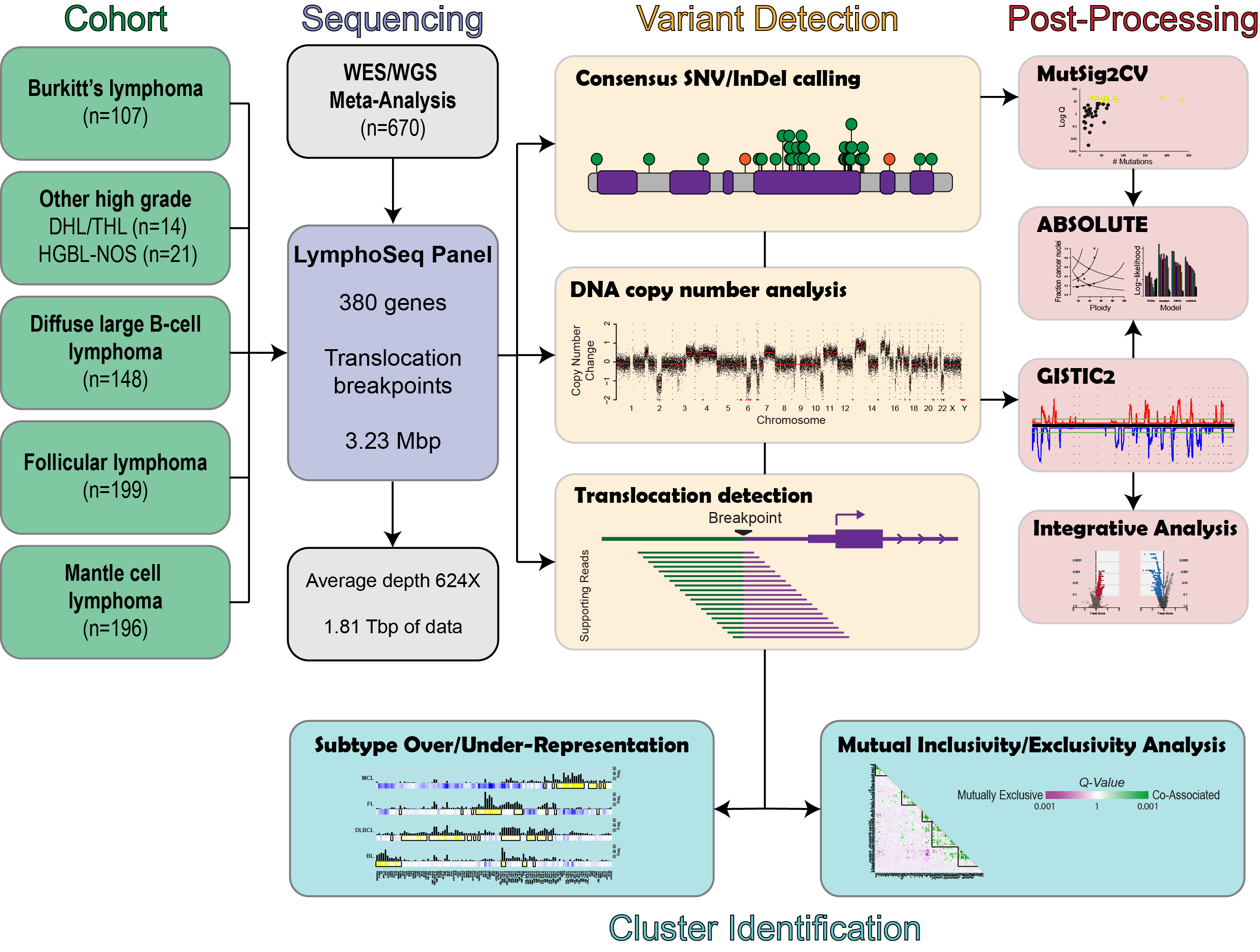

**Figure S1: A schematic overview of the study design.**

**Figure S2: Validation of CopyWriteR DNA copy number calls.** DNA copy number calculated from off-target NGS reads by CopyWriteR was compared to that calculated using Affymetrix 250K Nsp SNP microarrays for 3 FL tumors with available data. The copy number calls were highly concordant with the exception of 4 repetitive regions (arrows) that were identified as copy number losses by CopyWriteR, but not in the SNP microarray data. These regions were masked for the GISTIC2 analysis.

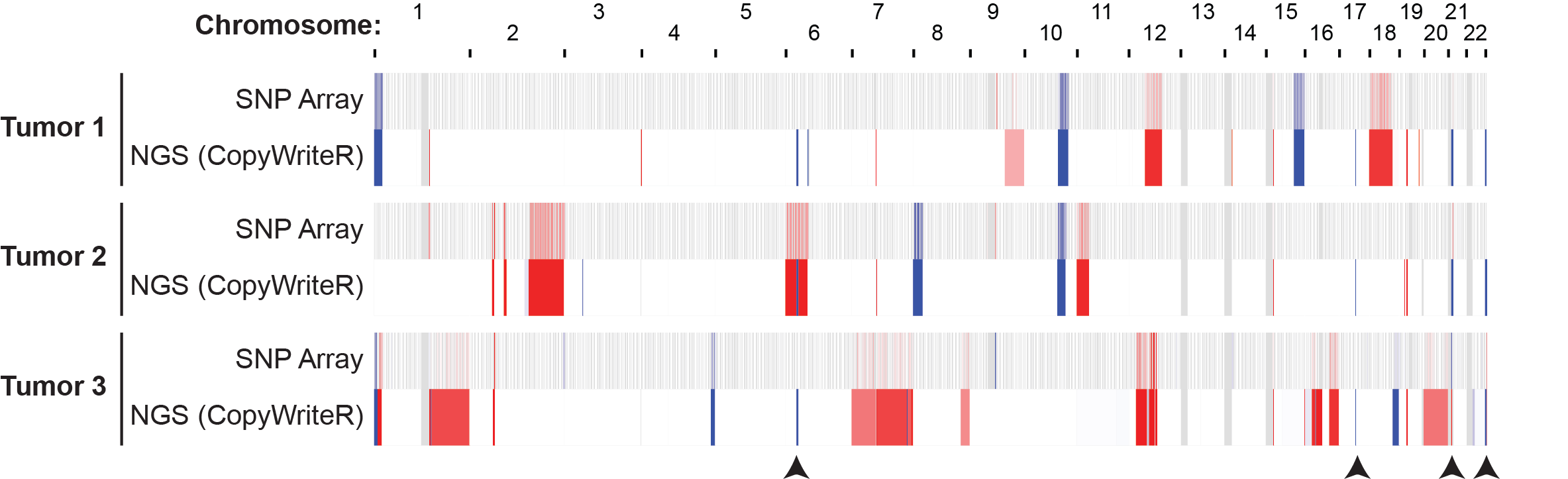

**Figure S3: Spectrum of mutational burden and copy number aberrant genome. A)** Mutational burden was calculated for the whole exome and from LymphoSeq targeted regions using previously published whole-exome data^11^, which showed a significant correlation. **B)** The mutational burden, AID-driven mutational burden, and percentage of copy number aberrant genome are shown for each tumor grouped by diagnosis. (*p<0.05, **p<0.01, ***p<0.001)

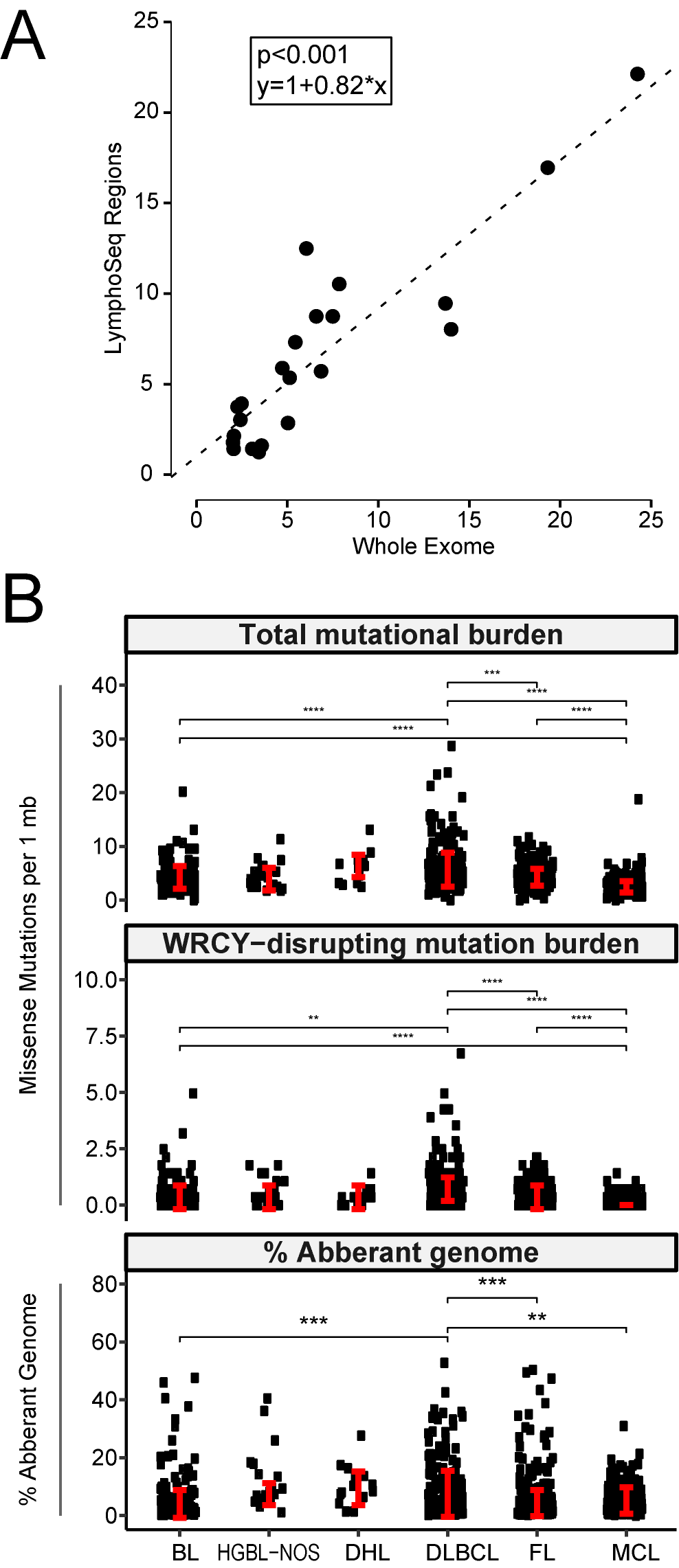

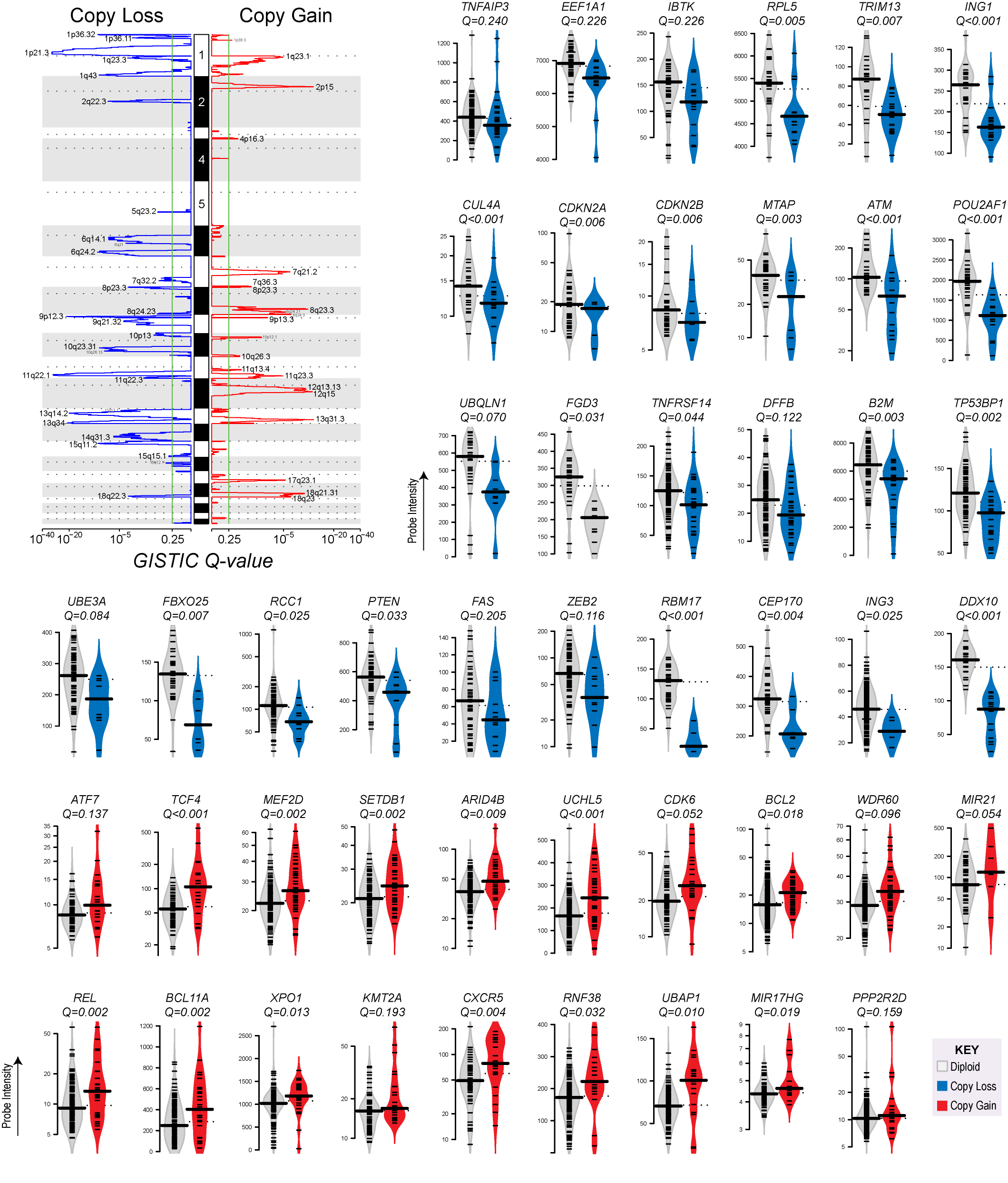
**Figure S4: GISTIC and integrative analysis.** A standard GISTIC plot for figure 2A is shown on the top left. Individual violin plots for the integrative analysis of genes appearing in figure 3 are also provided, colored by whether the gene is targeted by copy loss (blue) or copy gain (red).

**Figure S5: Comparison of DNA copy number alteration frequencies between NGS- and SNP microarray-based copy number analysis. A)** A heatmap shows the absolute DNA copy number of genes shown in Figure 3, as determined by SNP microarray analysis of 85 BL, 694 DLBCL, 404 FL and 206 MCL tumors. These same genes show frequent DNA copy number alterations in this independent cohort, as measured by SNP microarray. **B-E)** The frequencies of DNA copy number alterations for genes shown in Figure 3 are compared between NGS-based and SNP microarray-based DNA copy number analysis of independent cohorts. The frequencies significantly correlate in BL (B; Pearson’s correlation p-value < 0.0001, r = 0.8635), DLBCL (C; Pearson’s correlation p-value < 0.0001, r = 0.8519), FL (D; Pearson’s correlation p-value < 0.0001, r = 0.9233), and MCL (E; Pearson’s correlation p-value < 0.0001, r = 0.9683), providing validation of the NGS-based approach for DNA copy number analysis.

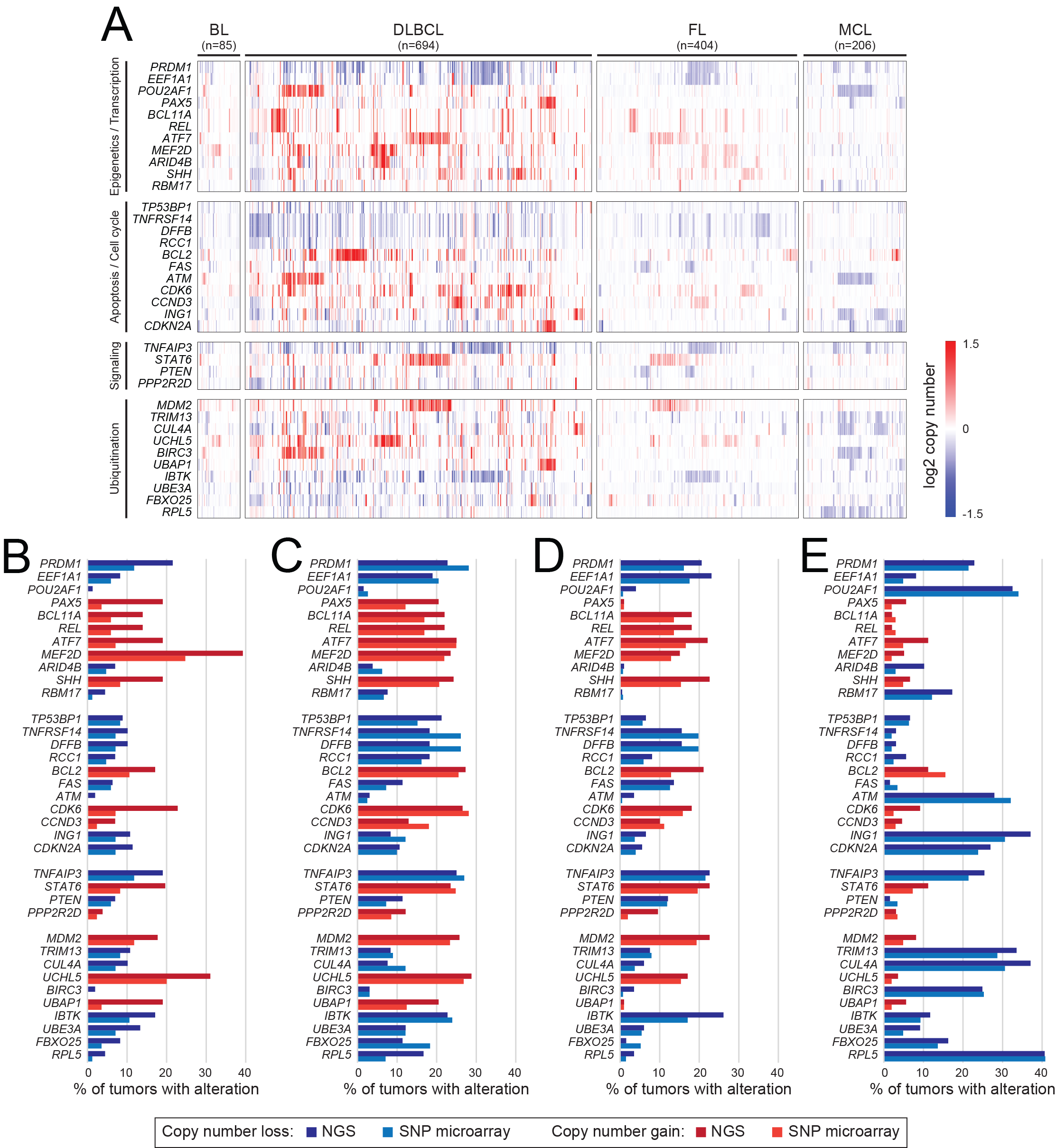

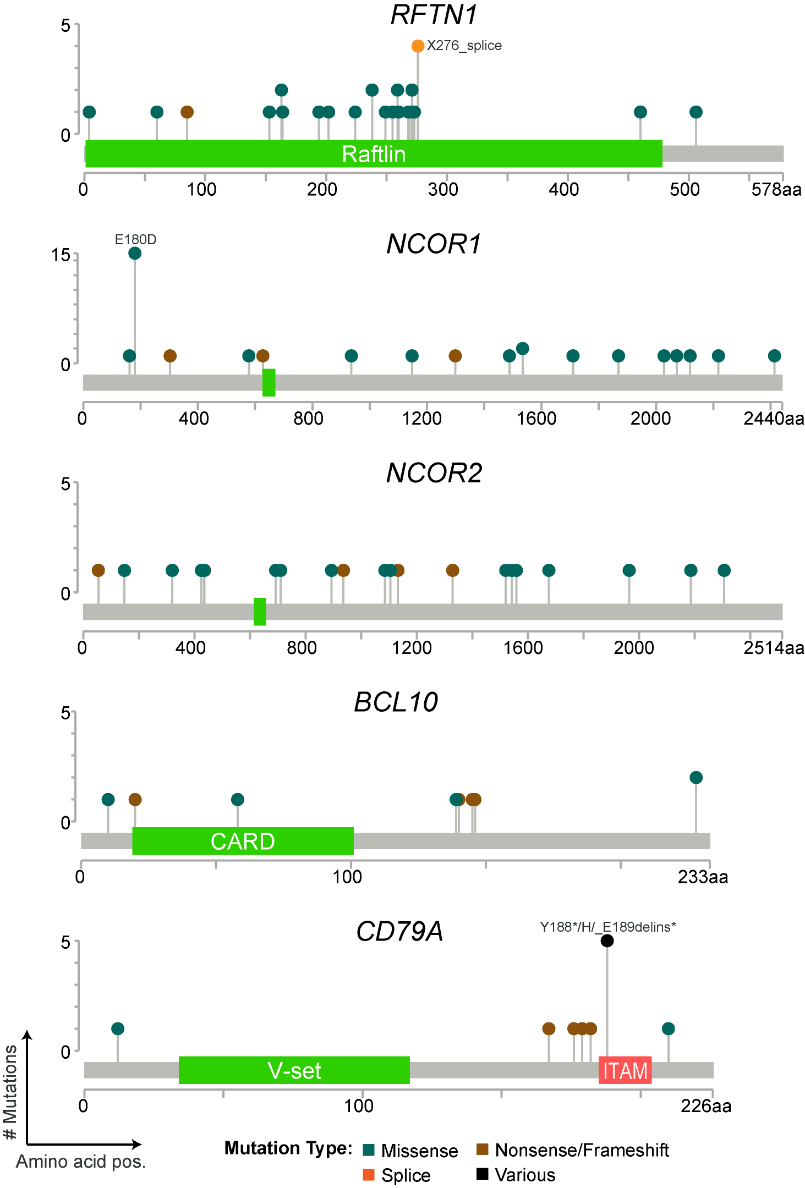
**Figure S6: Lollipop plots for select genes.**

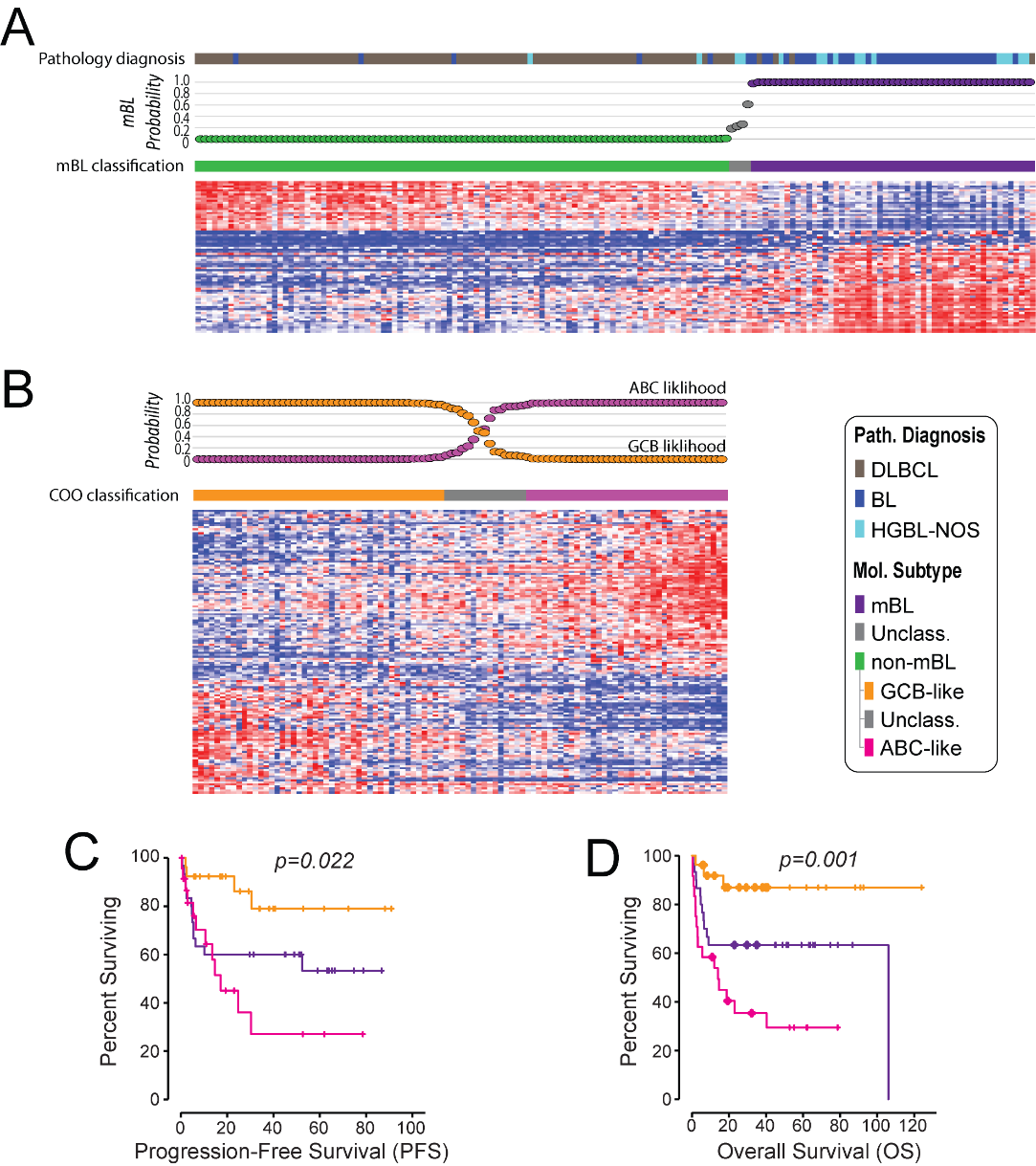

**Figure S7: Molecular classification of UNMC tumors. A)** Application of the molecular Burkitt lymphoma (mBL) Bayesian classifier to the tumors from the UNMC cohort. **B)** Application of the previously described cell of origin Bayesian classifier^41^ to the UNMC cohort. **C)** Progression-free survival of CHOP-treated patients according to their tumor’s molecular classification. **D)** Overall survival of CHOP-treated patients according to their tumor’s molecular classification.

**Supplementary References**

1	Khodadoust, M. S. *et al.* Antigen presentation profiling reveals recognition of lymphoma immunoglobulin neoantigens. *Nature* **543**, 723-727, doi:10.1038/nature21433 (2017).

2	Bea, S. *et al.* Landscape of somatic mutations and clonal evolution in mantle cell lymphoma. *Proceedings of the National Academy of Sciences of the United States of America* **110**, 18250-18255, doi:10.1073/pnas.1314608110 (2013).

3	Zhang, J. *et al.* The genomic landscape of mantle cell lymphoma is related to the epigenetically determined chromatin state of normal B cells. *Blood* **123**, 2988-2996, doi:10.1182/blood-2013-07-517177 (2014).

4	Schmitz, R. *et al.* Burkitt lymphoma pathogenesis and therapeutic targets from structural and functional genomics. *Nature* **490**, 116-120, doi:10.1038/nature11378 (2012).

5	Love, C. *et al.* The genetic landscape of mutations in Burkitt lymphoma. *Nature genetics* **44**, 1321-1325, doi:10.1038/ng.2468 (2012).

6	Morin, R. D. *et al.* Frequent mutation of histone-modifying genes in non-Hodgkin lymphoma. *Nature* **476**, 298-303, doi:10.1038/nature10351 (2011).

7	Morin, R. D. *et al.* Mutational and structural analysis of diffuse large B-cell lymphoma using whole-genome sequencing. *Blood* **122**, 1256-1265, doi:10.1182/blood-2013-02-483727 (2013).

8	Pasqualucci, L. *et al.* Analysis of the coding genome of diffuse large B-cell lymphoma. *Nature genetics* **43**, 830-837, doi:10.1038/ng.892 (2011).

9	Lohr, J. G. *et al.* Discovery and prioritization of somatic mutations in diffuse large B-cell lymphoma (DLBCL) by whole-exome sequencing. *Proceedings of the National Academy of Sciences of the United States of America* **109**, 3879-3884, doi:10.1073/pnas.1121343109 (2012).

10	Green, M. R. *et al.* Hierarchy in somatic mutations arising during genomic evolution and progression of follicular lymphoma. *Blood* **121**, 1604-1611, doi:10.1182/blood-2012-09-457283 (2013).

11	Green, M. R. *et al.* Mutations in early follicular lymphoma progenitors are associated with suppressed antigen presentation. *Proceedings of the National Academy of Sciences of the United States of America* **112**, E1116-1125, doi:10.1073/pnas.1501199112 (2015).

12	Okosun, J. *et al.* Integrated genomic analysis identifies recurrent mutations and evolution patterns driving the initiation and progression of follicular lymphoma. *Nature genetics* **46**, 176-181, doi:10.1038/ng.2856 (2014).

13	Pasqualucci, L. *et al.* Genetics of follicular lymphoma transformation. *Cell reports* **6**, 130-140, doi:10.1016/j.celrep.2013.12.027 (2014).

14	Landau, D. A. *et al.* Evolution and impact of subclonal mutations in chronic lymphocytic leukemia. *Cell* **152**, 714-726, doi:10.1016/j.cell.2013.01.019 (2013).

15	Puente, X. S. *et al.* Whole-genome sequencing identifies recurrent mutations in chronic lymphocytic leukaemia. *Nature* **475**, 101-105, doi:10.1038/nature10113 (2011).

16	Quesada, V. *et al.* Exome sequencing identifies recurrent mutations of the splicing factor SF3B1 gene in chronic lymphocytic leukemia. *Nature genetics* **44**, 47-52, doi:10.1038/ng.1032 (2012).

17	Wang, L. *et al.* SF3B1 and other novel cancer genes in chronic lymphocytic leukemia. *The New England journal of medicine* **365**, 2497-2506, doi:10.1056/NEJMoa1109016 (2011).

18	Kiel, M. J. *et al.* Whole-genome sequencing identifies recurrent somatic NOTCH2 mutations in splenic marginal zone lymphoma. *The Journal of experimental medicine* **209**, 1553-1565, doi:10.1084/jem.20120910 (2012).

19	Rossi, D. *et al.* The coding genome of splenic marginal zone lymphoma: activation of NOTCH2 and other pathways regulating marginal zone development. *The Journal of experimental medicine* **209**, 1537-1551, doi:10.1084/jem.20120904 (2012).

20	Chapman, M. A. *et al.* Initial genome sequencing and analysis of multiple myeloma. *Nature* **471**, 467-472, doi:10.1038/nature09837 (2011).

21	Treon, S. P. *et al.* MYD88 L265P somatic mutation in Waldenstrom's macroglobulinemia. *The New England journal of medicine* **367**, 826-833, doi:10.1056/NEJMoa1200710 (2012).

22	Gunawardana, J. *et al.* Recurrent somatic mutations of PTPN1 in primary mediastinal B cell lymphoma and Hodgkin lymphoma. *Nature genetics* **46**, 329-335, doi:10.1038/ng.2900 (2014).

23	Vater, I. *et al.* The mutational pattern of primary lymphoma of the central nervous system determined by whole-exome sequencing. *Leukemia* **29**, 677-685, doi:10.1038/leu.2014.264 (2015).

24	Li, H. Aligning sequence reads, clone sequences and assembly contigs with BWA-MEM. *arXiv* **1303.3997** (2013).

25	McKenna, A. *et al.* The Genome Analysis Toolkit: a MapReduce framework for analyzing next-generation DNA sequencing data. *Genome research* **20**, 1297-1303, doi:10.1101/gr.107524.110 (2010).

26	Ng, S. B. *et al.* Targeted capture and massively parallel sequencing of 12 human exomes. *Nature* **461**, 272-276, doi:10.1038/nature08250 (2009).

27	Lek, M. *et al.* Analysis of protein-coding genetic variation in 60,706 humans. *Nature* **536**, 285-291, doi:10.1038/nature19057 (2016).

28	Lawrence, M. S. *et al.* Mutational heterogeneity in cancer and the search for new cancer-associated genes. *Nature* **499**, 214-218, doi:10.1038/nature12213 (2013).

29	Kuilman, T. *et al.* CopywriteR: DNA copy number detection from off-target sequence data. *Genome biology* **16**, 49, doi:10.1186/s13059-015-0617-1 (2015).

30	Mermel, C. H. *et al.* GISTIC2.0 facilitates sensitive and confident localization of the targets of focal somatic copy-number alteration in human cancers. *Genome biology* **12**, R41, doi:10.1186/gb-2011-12-4-r41 (2011).

31	Reich, M. *et al.* GenePattern 2.0. *Nature genetics* **38**, 500-501, doi:10.1038/ng0506-500 (2006).

32	Bouska, A. *et al.* Adult high-grade B-cell lymphoma with Burkitt lymphoma signature: genomic features and potential therapeutic targets. *Blood* **130**, 1819-1831, doi:10.1182/blood-2017-02-767335 (2017).

33	Green, M. R. *et al.* Transient expression of Bcl6 is sufficient for oncogenic function and induction of mature B-cell lymphoma. *Nature communications* **5**, 3904, doi:10.1038/ncomms4904 (2014).

34	Li, M. *et al.* Non-oncogene Addiction to SIRT3 Plays a Critical Role in Lymphomagenesis. *Cancer cell* **35**, 916-931 e919, doi:10.1016/j.ccell.2019.05.002 (2019).

35	Parsa, S. *et al.* The serine hydroxymethyltransferase-2 (SHMT2) initiates lymphoma development through epigenetic tumor suppressor silencing. *Nature Cancer* **1**, 653-664, doi:10.1038/s43018-020-0080-0 (2020).

36	Huang, D. W. *et al.* DAVID Bioinformatics Resources: expanded annotation database and novel algorithms to better extract biology from large gene lists. *Nucleic acids research* **35**, W169-175, doi:10.1093/nar/gkm415 (2007).

37	Lenz, G. *et al.* Stromal gene signatures in large-B-cell lymphomas. *The New England journal of medicine* **359**, 2313-2323, doi:10.1056/NEJMoa0802885 (2008).

38	Dave, S. S. *et al.* Molecular diagnosis of Burkitt's lymphoma. *The New England journal of medicine* **354**, 2431-2442, doi:10.1056/NEJMoa055759 (2006).

39	Iqbal, J. *et al.* Genome-wide miRNA profiling of mantle cell lymphoma reveals a distinct subgroup with poor prognosis. *Blood* **119**, 4939-4948, doi:10.1182/blood-2011-07-370122 (2012).

40	Hummel, M. *et al.* A biologic definition of Burkitt's lymphoma from transcriptional and genomic profiling. *The New England journal of medicine* **354**, 2419-2430, doi:10.1056/NEJMoa055351 (2006).

41	Wright, G. *et al.* A gene expression-based method to diagnose clinically distinct subgroups of diffuse large B cell lymphoma. *Proceedings of the National Academy of Sciences of the United States of America* **100**, 9991-9996, doi:10.1073/pnas.1732008100 (2003).
